# The role of V3 neurons in speed-dependent interlimb coordination during locomotion in mice

**DOI:** 10.1101/2021.09.01.458603

**Authors:** Han Zhang, Natalia A. Shevtsova, Dylan Deska-Gauthier, Colin Mackay, Kimberly J. Dougherty, Simon M. Danner, Ying Zhang, Ilya A. Rybak

**Affiliations:** Department of Medical Neuroscience, Brain Repair Centre, Faculty of Medicine, Dalhousie University, Halifax, NS, Canada; Department of Neurobiology and Anatomy, College of Medicine, Drexel University, Philadelphia, PA, United States

## Abstract

Speed-dependent interlimb coordination allows animals to maintain stable locomotion under different circumstances. We have previously demonstrated that a subset of spinal V3 neurons contributes to stable locomotion by mediating mutual excitation between left and right lumbar rhythm generators (RGs). Here, we expanded our investigation to the V3 neurons involved in ascending long propriospinal interactions (aLPNs). Using retrograde tracing, we revealed a subpopulation of lumbar V3 aLPNs with contralateral cervical projections. V3^OFF^ mice, in which all V3 neurons were silenced, had a significantly reduced maximal locomotor speed, were unable to move using stable trot, gallop, or bound, and predominantly used lateral-sequence walk. To understand the functional roles of V3 aLPNs, we adapted our previous model of spinal circuitry controlling quadrupedal locomotion (Danner et al., 2017), by incorporating diagonal V3 aLPNs mediating inputs from each lumbar RG to the contralateral cervical RG. The updated model reproduces our experimental results and suggests that locally projecting V3 neurons, mediating left–right interactions within lumbar and cervical cords, promote left–right synchronization necessary for gallop and bound, whereas the V3 aLPNs promote synchronization between diagonal fore and hind RGs necessary for trot. The model proposes the organization of spinal circuits available for future experimental testing.

## Introduction

Coordinated rhythmic movement of the limbs during locomotion in mammals is primarily controlled by neural circuitry within the spinal cord. This spinal circuitry includes rhythm-generating circuits (Graham Brown, 1911, 1914) and multiple commissural, propriospinal, premotor, and pattern formation neurons (Grillner, 2006; Kiehn, 2006, 2011, 2016; Rybak et al., 2006a, 2006b, 2013, 2015; Jankowska, 2008; McCrea and Rybak, 2008; Danner et al., 2016, 2017, 2019; Ausborn et al., 2021). It is commonly accepted that each limb is controlled by a separate spinal rhythm generator (RG) (Forssberg et al., 1980; Thibaudier et al., 2013; Frigon, 2017; Danner et al., 2019; Latash et al., 2020) and that RGs controlling left and right forelimbs and left and right hindlimbs are located on the corresponding sides of cervical and lumbar enlargements of the spinal cord, respectively (Kato, 1990; Ballion et al., 2001; Juvin et al., 2005, 2012). The left and right lumbar and cervical circuits are connected through multiple types of local commissural interneurons (CINs), which coordinate left–right activities (Stein, 1976; Butt and Kiehn, 2003; Quinlan and Kiehn, 2007; Talpalar et al., 2013; Bellardita and Kiehn, 2015; Rybak et al., 2015; Shevtsova et al., 2015). In turn, descending and ascending long propriospinal neurons (LPNs) mediate interactions between cervical and lumbar circuits (Juvin et al., 2005; Dutton et al., 2006; Reed et al., 2006; Brockett et al., 2013; Ruder et al., 2016; Flynn et al., 2017; Pocratsky et al., 2020). Diverse populations of CINs and LPNs are involved in coordination of limb movements, defining locomotor gait, and controlling speed and balance during locomotion (Danner et al., 2016, 2017; Kiehn, 2016; Ruder et al., 2016; Pocratsky et al., 2020).

Experimental studies with genetic ablation, silencing, or activation of genetically identified neuron types, such as V0_V_, V0_D_, V1, V2a, V2b, V3, Shox2, and Hb9, allowed partial identification and/or suggestion of neuron-type specific roles in spinal circuits and motor control, including locomotion (Lanuza et al., 2004; Gosgnach et al., 2006; Crone et al., 2008; Zhang et al., 2008; Goulding, 2009, 2009; Dougherty et al., 2013; Talpalar et al., 2013; Shevtsova et al., 2015; Bikoff et al., 2016; Kiehn, 2016; Caldeira et al., 2017; Ziskind-Conhaim and Hochman, 2017; Dougherty and Ha, 2019; Falgairolle and O’Donovan, 2019). However, so far these studies have mostly focused on lumbar circuits. Genetic identities have only begun to be ascribed to long propriospinal pathways connecting lumbar and cervical spinal segments (Ruder et al., 2016; Flynn et al., 2017). Removal of descending LPNs (dLPNs) resulted in transient periods of disordered left–right coordination in mice (Ruder et al., 2016); an effect that our previous computational model (Danner et al., 2017) attributed to excitatory diagonally-projecting (commissural) V0_V_ LPNs. Similarly, more recent experimental silencing of ascending LPNs (aLPNs) was shown to affect left–right coordination in rats in certain locomotor contexts (Pocratsky et al., 2020); however, the specific populations involved in the effects are unknown. Although the excitatory V3 neurons are unlikely to have significant descending propriospinal projections (Ruder et al., 2016; Flynn et al., 2017), it remains unknown whether there is a subpopulation of V3 neurons that are aLPNs with involvement in fore–hind and/or left–right interlimb coordination.

In the present study, we specifically focused on the potential role of V3 neurons in long propriospinal interactions between lumbar and cervical circuits controlling interlimb coordination. These neurons are defined by post-mitotic expression of the transcription factor single-minded homolog 1 (Sim1). They are excitatory neurons and the majority of them project to the contralateral side of the spinal cord (Zhang et al., 2008). Results from initial V3 silencing experiments suggested that V3 neurons are involved in the control of locomotion, specifically robustness of the rhythm and left–right coordination (Zhang et al., 2008). Our prior modeling studies suggest that V3 commissural neurons are involved in promoting left–right synchronization during synchronous gaits (Rybak et al., 2013, 2015; Shevtsova et al., 2015; Danner et al., 2016, 2017; Ausborn et al., 2021) by providing mutual excitation between the extensor half-centers of the left and right lumbar RGs (Danner et al., 2019). Yet, these studies did not provide an explanation for the unbalanced locomotion and variable left–right coordination after inactivation of V3 neurons (Zhang et al., 2008). Also, these studies did not account for heterogeneity of the V3 population, which was shown to contain distinct subpopulations with different biophysical properties, laminar distributions, and connectivity (Borowska et al., 2013, 2015; Blacklaws et al., 2015; Chopek et al., 2018), which may underly different functions.

Here, we identified a subset of V3 neurons with cell bodies in the lumbar spinal cord that have direct excitatory projections to the contralateral side of the cervical enlargement. The recruitment of these neurons increased with locomotor speed. We also show that mice with glutamatergic transmission conditionally knocked-out in V3 neurons (V3^OFF^ mice) had significantly reduced maximal speeds of locomotion. Moreover, even moving with relatively low and medium speeds, V3^OFF^ mice lost the ability to trot stably, replacing this most typical mouse gait with a lateral-sequence walk. At higher locomotor speeds, V3^OFF^ mice exhibited high step-to-step variability of left–right coordination, which could be a reason for the speed limitation observed in these animals. To determine potential connectivity and functions of local and long propriospinal V3 neurons in spinal locomotor circuits, we updated and extended our previous computational model of spinal circuits consisting of four RGs coupled by multiple CIN and LPN pathways (Danner et al., 2017). We included V3 aLPNs providing diagonal RG synchronization necessary for trot in addition to the local V3 CINs involved in left–right synchronization necessary for gallop and bound. The updated model reproduced speed-dependent gait expression in wild type (WT) mice as well as the experimentally detected changes in the maximal speed, interlimb coordination, and gait expression in V3^OFF^ mice. The model predicts possible interactions between the V3 aLPNs and some local CINs. Taken together, our results suggest different functional roles of the local V3 CINs and V3 aLPNs in interlimb coordination and speed-dependent gait expression during locomotion.

## Results

### Experimental studies *in vitro* and *in vivo*

#### Lumbar propriospinal V3 interneurons provide ascending excitatory drives to the contralateral cervical locomotor circuits

Several subpopulations of V3 neurons have been found and characterized in the mouse spinal cord (Borowska et al., 2013; Chopek et al., 2018; Deska-Gauthier et al., 2020). Until now, however, it has not been shown whether any V3 neurons can serve as long propriospinal neurons (LPNs) to connect the spinal circuits in lumbar and cervical regions for fore–hind coordination. Previous studies indicated that there might be only a limited number of excitatory descending LPNs (dLPNs) projecting from cervical to lumbar regions (Ruder et al., 2016; Flynn et al., 2017). Therefore, we primarily focused on studying the ascending LPNs (aLPNs) with projections from lumbar to cervical regions for potential overlap with the V3 neuronal population. To do so, we injected a retrograde tracer, cholera-toxin B (CTB), into the cervical C5 to C8 region of Sim1tdTomato mice (Figure 1A1). After 7 days, we harvested the lumbar spinal cords, and then identified and mapped the tdTomato (tdTom) fluorescent protein and CTB double positive neurons in lumbar cross-sections (Figure 1A2). CTB and tdTom positive neurons should mainly be of V3 type with ascending projections to the cervical region (V3 aLPNs). We found that these V3 aLPNs were almost exclusively located contralaterally to the cervical injection sites. Overall, 30% of V3 neurons (n=3 mice) in L1 to L3 were stained by CTB but only 12% of V3 neurons in L4 to L6 were CTB positive. Even though clusters of V3 neurons were distributed across ventral to deep dorsal horns (Zhang et al., 2008; Borowska et al., 2013, 2015; Blacklaws et al., 2015), the highest density of V3 aLPNs was found in deep dorsal horn, lamina IV to VI, in rostral lumbar segments (Figure 1A3). A relatively small number of these neurons were located in the intermediate and ventral regions, lamina VII and VIII, in caudal lumbar segments (Figure 1A4).

**Figure 1.**
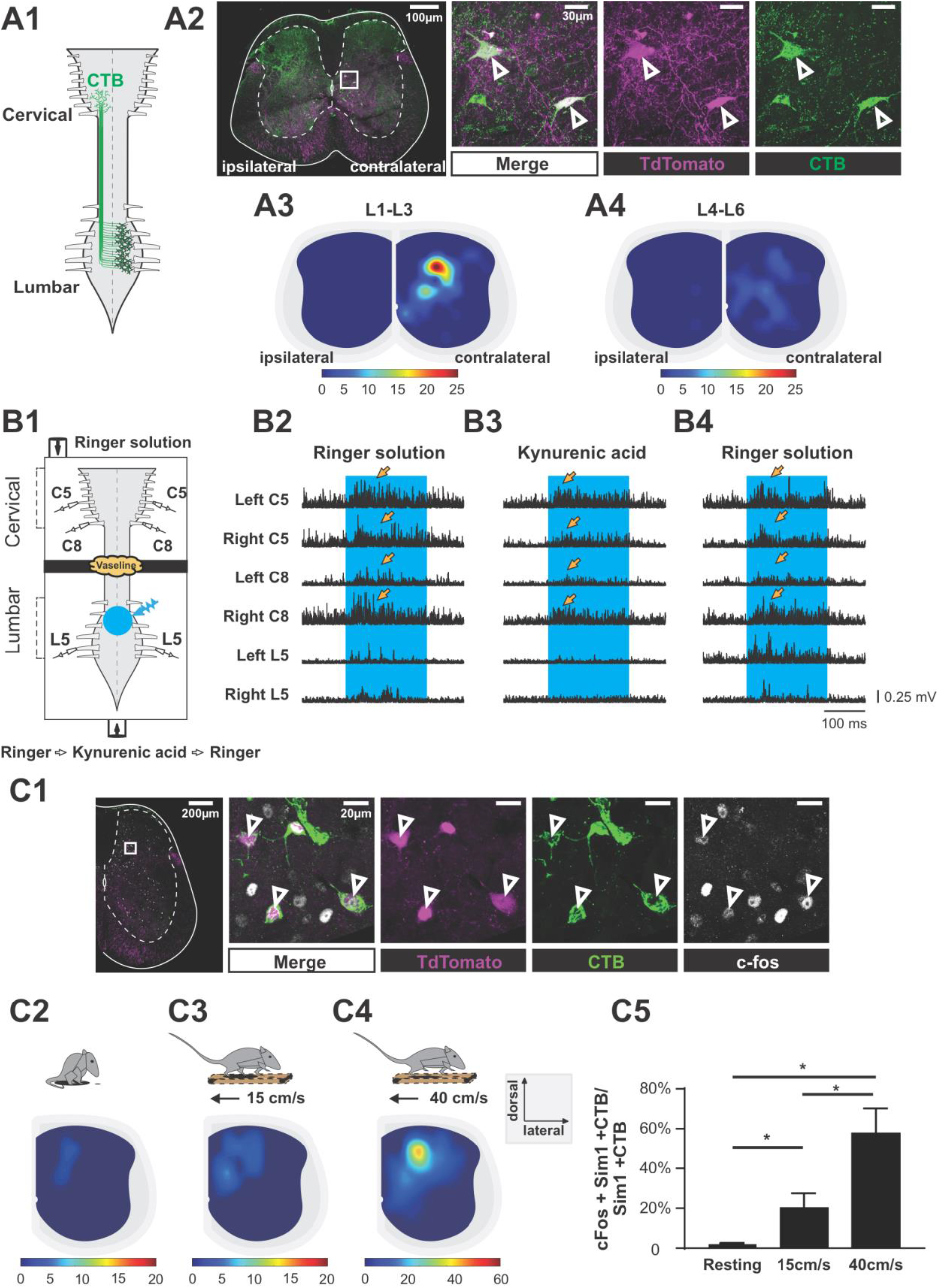
Ascending V3 long propriospinal neurons identified in lumbar spinal cord. **A1.** Illustration of the experimental strategy to identify lumbar ascending V3 LPNs. CTB (green) injected into the cervical region is picked up at axon terminals and retrogradely transported to cell bodies located in the lumbar region. **A2.** Representative image of cross section of the lumbar spinal cord of Sim1Cre/+; Rosafloxstop26TdTom mouse (far left panel). The ‘ipsilateral’ and ‘contralateral’ side are relative to the injection side in the cervical region. Immunohistochemical staining with different antibodies to illustrate the neurons expressing TdTom (magenta) and CTB (green) (right panels); **A3** and **A4.** Colour coded heat maps of the distribution pattern of ascending V3 LPNs in the rostral lumbar (**A3**) and caudal lumbar (**A4**) segments. Scale bars of cell numbers for heat maps are shown below. **B1**. Schematic of the experimental strategy. Isolated neonatal spinal cord with ventral roots attached is placed in a split bath recording chamber. The chamber is partitioned into two sides with a Vaseline wall (represented by yellow cloud in the figure). The ENG activities of left and right cervical lumbar ventral roots were recorded. The blue-fluorescent light indicated by the blue transparent circle is on the ventral side of rostral lumbar, L1-L3, segments; **B2-B4**. representative traces of rectified ENG recordings at ventral roots of both sides of cervical (C5 and C8) and L5 lumbar spinal segments of P2 SimAi32 mouse, before (B2), during (B3), and after wash-out (B4) of kynurenic acid application. The time period of optical stimulation is indicated with the blue shadowed area. The persistent cervical ventral root activities responding to lumbar optical stimulation under all conditions are indicated with yellow arrows. **C1**. Representative image of TdTom+ V3 INs (magenta), CTB (green) and c-Fos (white) of a half cross-section of the spinal cord from a Sim1TdTom mouse. Enlarged images showing triple positive cells. (scale bars = 200μM and 20μM); **C2-C4**. Colour-coded heat maps showing the distribution of c-Fos, CTB and TdTom triple-positive V3 INs on the cross section of lumbar spinal cord during resting (**C2**), 15 cm/s (**C3**) and 40 cm/s (**C4**) treadmill locomotion (nresting=3, n15cm/s=3, n40cm/s =3); 5, Histogram of the corresponding percentage of triple-positive cells in TdTom/CTB double positive population in L1 to L3 segments. *, 0.01 < P < 0.05.

To test whether these ascending V3 LPNs in the lumbar segment affect locomotor circuits and motor outputs in the cervical region, we employed Sim1^Cre/+^; Ai32 (Sim1Ai32) mice, which expressed channelrhodopsin2 (ChR2) specifically in Sim1 positive V3 neurons. Isolated spinal cords from P2-3 Sim1Ai32 mice were placed in a perfusion chamber split into two compartments. This split chamber was constructed over the thoracic T6-T8 segments with petroleum jelly (Vaseline) walls (Fig 1B1). Suction electrodes for electroneurogram (ENG) recording were placed on the lumbar and cervical ventral roots (Figure 1B1). We found that photo-activation of V3 neurons in the lumbar region evoked strong activity in all recorded lumbar and cervical ventral roots (Figure 1B2). Then, to test whether this excitation measured in the cervical roots was provided directly by ascending projections of lumbar V3 neurons, we blocked the glutamatergic transmission selectively in the lumbar region with 2 mM of kynurenic acid (KYN), which completely blocks NMDA and AMPA/kainate receptors (Hägglund et al., 2010). We found that optical stimulation of lumbar V3 neurons did not evoke any response in lumbar ventral roots, but the motor responses from the cervical roots were still present (Figure 1B3). The lumbar responses reappeared after the drug was washed out (Figure 1B4). These results demonstrate that lumbar V3 neurons directly innervate neurons in cervical motor circuits.

Next, we tested whether V3 aLPNs were active during locomotion. We injected CTB in the cervical region of young adult (P35–40) Sim1tdTom mice. Seven days post injection, we subjected the animals to treadmill locomotion at either 15 cm/s or 40 cm/s. The control group was sitting in the cage for one hour. After one-hour of rest, which maximizes c-Fos expression in neurons (Dai et al., 2005), we harvested the lumbar spinal cord and used immunohistochemical staining to detect the expression of c-Fos protein in V3 neurons (Figure 1C1; <u>Blacklaws et al., 2015</u>). We found that the number of triple labelled (c-Fos/CTB/tdTom) V3 neurons significantly increased after treadmill locomotion at both speeds compared to animals that only rested, but the percentage of these triple positive V3 aLPNs almost tripled at 40 cm/s compared to 15 cm/s (Figure 1C2–C5).

Taken together, we conclude that there are subsets of lumbo-cervical projecting commissural V3 neurons providing excitatory drive to cervical locomotor networks, particularly at medium locomotor speeds.

#### Elimination of V3 neurons in the mouse spinal cord limits the locomotor speed

To study the contribution of V3 neurons to the control of locomotion and interlimb coordination, we generated Sim1^Cre/+^;VGluT2^flox/flox^ (V3^OFF^) mice, in which the expression of Vesicular Glutamate Transporter 2 (vGluT2) was deleted in V3 neurons (Chopek et al., 2018). We then subjected the control (WT) and V3^OFF^ mice to a treadmill locomotion at different speeds. We found that the highest speed that the V3^OFF^ mice could reach was around 35 cm/s (mean = 34.17 ± 6.69 cm/s; Figure 2), and only 3 out of 11 mice could reach 40 cm/s (the maximal speed included for the subsequent analysis). Under the same experimental conditions, however, WT mice could run with speeds up to 75 ± 7.56 cm/s (Figure 2). This result indicates that V3 neurons are essential for high-speed locomotion in mice.

**Figure 2.**
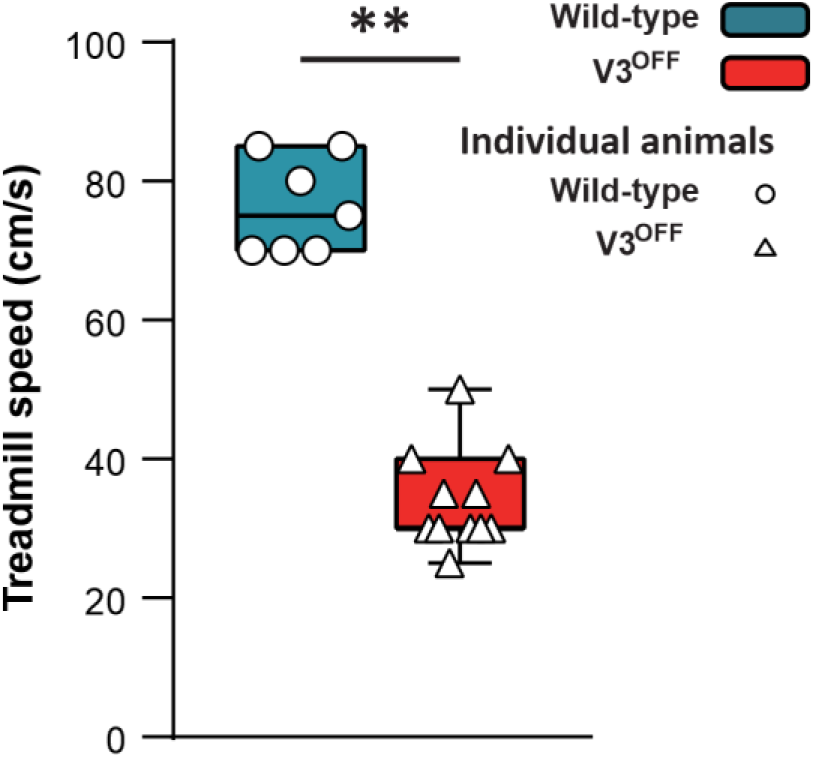
Maximum speeds of WT and V3^OFF^ mice on the treadmill. The box and whisker plots showing the highest speed of the individual WT mice (n=7) and V3^OFF^ mice (n=11) on the treadmill. **, 0.001 < P < 0.01

#### Elimination of V3 neurons changed speed-dependent interlimb coordination

To understand the changes in interlimb coordination caused by V3 silencing at low and medium speeds, we calculated the phase differences (relative phases) between different pairs of limbs in both WT and V3^OFF^ mice: homologous (left–right), between two forelimbs and two hindlimbs; homolateral, between left forelimbs and hindlimbs; and diagonal, between left hindlimb and right forelimb (Figure 3 and Figure 3—figure supplement 1). We then compared the corresponding relative phases and their variability between WT and V3^OFF^ mice on average (Figure 3A1–D1) and at different treadmill speeds (Figure 3A2–D2, Figure 3—figure supplement 1).

**Figure 3.**
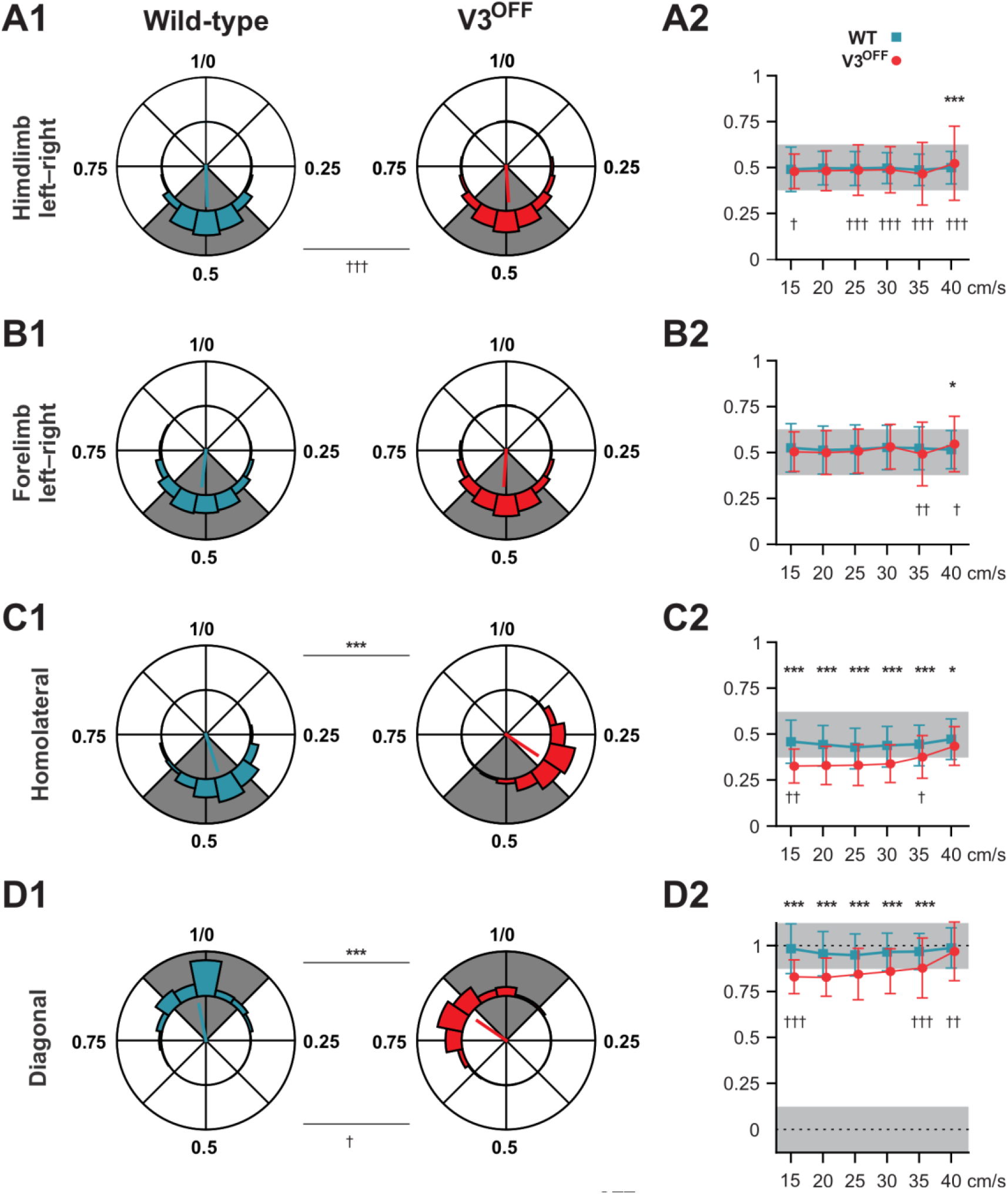
Interlimb coordination in WT and V3^OFF^ mice at different speeds. **A1-D1.** Circular plots of hindlimb (**A1**) and forelimb (**B1**) left–right phase differences, homolateral phase differences (**C1**) and diagonal phase differences (**D1**) in wild type (WT; blue) and V3^OFF^ (red) mice. Except for the forelimb left–right phase differences, the left hindlimbs are used as the reference limb. Each vector, blue line (WT) and red line (V3^OFF^), in the circular plot, indicates the mean value (direction) and robustness (radial line/length) of the phase differences. The circle is evenly separated into 8 fractions. The circular histograms represent the distribution of phase differences of all steps at all tested speeds (WT n = 1292; V3^OFF^ n = 1478). **A2-D2**. Plots of mean values of coupling phases at individual speeds of V3^OFF^ (red) and WT (blue) mice. *, P < 0.01; **, P < 0.001; ***, P < 0.0001 for comparisons of mean phase differences; †, P < 0.01; ††, P < 0.001; †††, P < 0.0001 for comparisons of the variability (concentration parameter κ) of the phase differences.

The left–right hindlimb and forelimb phase differences did not significantly differ between WT and V3^OFF^ mice across speeds (Figure 3A1,B1,A2,B2, Figure 3—figure supplement 1 A,B); mean values were close to 0.5 (perfect alternation) in both WT and V3^OFF^ mice. The tests at individual treadmill speeds showed no significant differences, except at 40 cm/s, where the mean forelimb and hindlimb left–right phase differences of V3^OFF^ mice differed slightly from those of the WT mice (Figure 3—figure supplement 1A,B). However, the variability of left–right phase differences in V3^OFF^ mice significantly increased compared to WT mice at higher locomotor speeds (Figure 3A2,B2).

In contrast to the left–right phase differences, the mean values of the homolateral (Figure 3C1,C2, Figure 3—figure supplement 1C) and diagonal phase differences (Figure 3D1,D2, Figure 3—figure supplement 1D) in V3^OFF^ mice significantly deviated from those in WT mice. Tests for individual speeds showed that these differences were significant across almost all tested speeds. At the highest speed (40 cm/s), the difference of the homolateral phase-differences was smaller than that at lower speeds but still significant, and the diagonal phase-differences did not differ significantly. This indicates that, at least at low to medium speeds, the V3^OFF^ mice were unable to support diagonal synchronization and homolateral alternation.

#### Elimination of V3 neurons changes the preferred gait from trot to walk at intermediate speeds and causes gait instability at higher speeds

Phase relationships between the four limbs define locomotor gaits (Hildebrand, 1976, 1980, 1989; Bellardita and Kiehn, 2015; Lemieux et al., 2016). Thus, the changes in the homolateral and diagonal phasing after functional removal of V3 neurons should impact gait expression.

Indeed, the step patterns and gait expression of V3^OFF^ mice were altered in a speed-dependent manner compared to WT litter mates. Figure 4B1 and B2 shows representative stance phases of all four limbs at different treadmill speeds in WT and V3^OFF^ mice, respectively.

**Figure 4.**
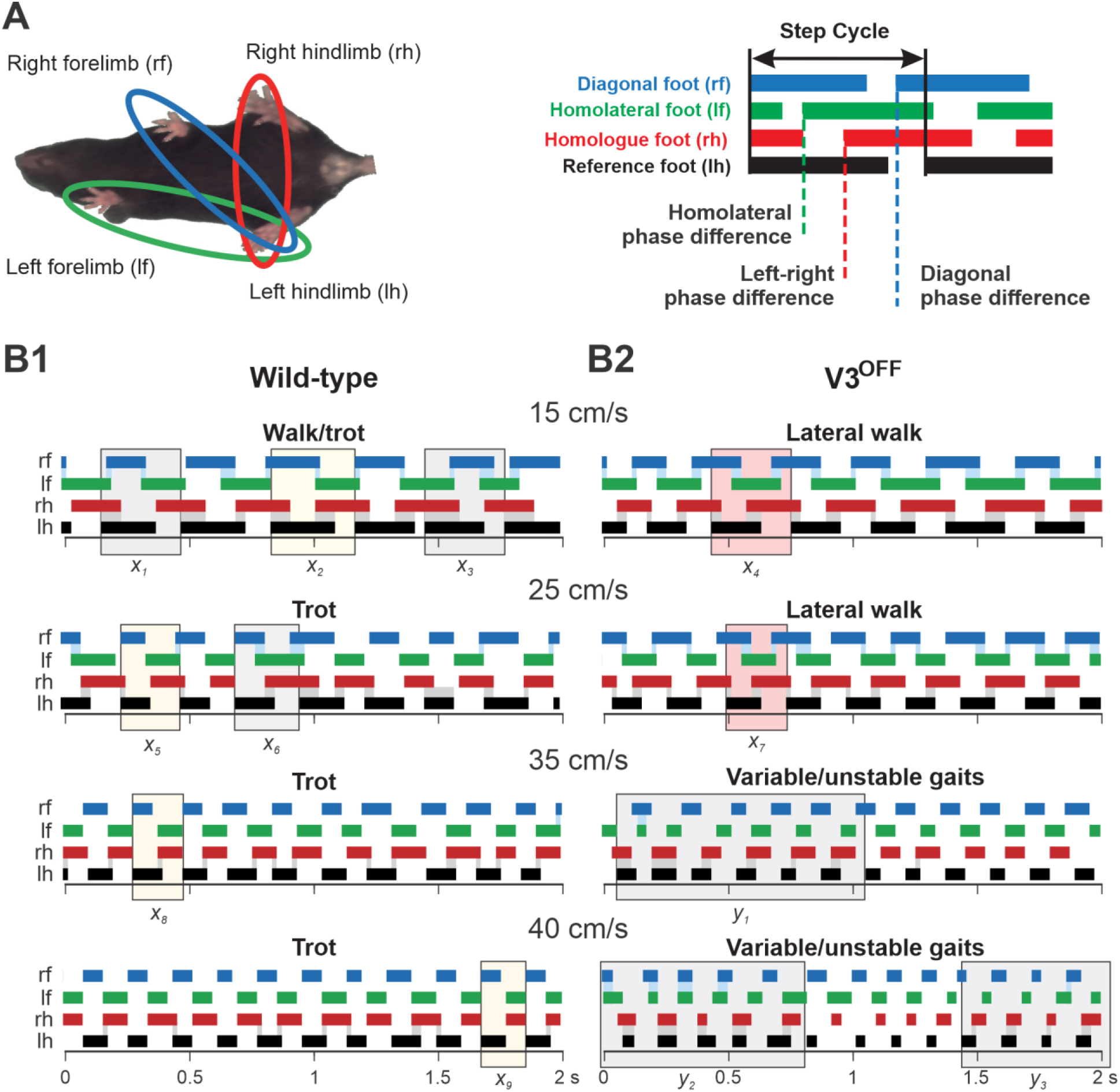
Step patterns of WT and V3OFF mice at different speeds. **A.** Illustration of limb-couplings (left). The footprint diagrams of individual limb are represented by color-coded bar graphs (right): the stance phase of each step is shown with solid bar and swing phase is the interval between two bars. The step cycle is measured from the duration between the onset of contacts of two consecutive steps of the same foot. Phase differences between the limbs was calculated as the interval between the start of stance of a specific limb and the reference limb divided by the period or step-cycle duration of the reference limb. **B1-B2.** Representative stance phases of 2 s episodes of WT (**B1**) and V3^OFF^ (**B2**) mice at low (15 cm/s and 25 cm/s) and medium (35cm/s and 40cm/s) treadmill speeds. *x*_1–9_, exemplary steps; *y*_1–3_, exemplary episodes referred to from the text. The shading colour indicates the gait (red, lateral-sequence walk; yellow, trot; grey, undefined). The black bars show the stance phase of the reference foot (left hindlimb). Lateral walk: lateral-sequence walk.

At low treadmill speeds (15 and 25 cm/s), WT mice mainly used trot (a two-beat gait characterized by diagonal synchronization and left–right alternation; Figure 4B1, e.g., x_2_, x_5_); however, these are transition speeds and the WT mouse exhibited a comparatively high step-to-step variability with steps that could be classified as different types of walks or even canter (a three-beat gait with only one diagonal synchronized) interspersed (e.g., steps *x*_1_, *x*_3_, *x*_6_) between trot steps. At the same low treadmill speeds (15 and 25 cm/s), V3^OFF^ mice used a lateral-sequence walk (a four-beat gait with longer stance than swing phases where the stance of a hindlimb is followed by that of the ipsilateral forelimb; Figure 4B2: e.g., steps *x*_4_, *x*_7_). The lateral-sequence walk of V3^OFF^ mice was very stable and consistent with very little variability between the different step-cycles (Figure 4B2).

At higher treadmill speeds (35 and 40 cm/s), the step pattern of V3^OFF^ mice became unstable (Figure 4B2), while the WT mice exhibited consistent trot steps with little step-to-step variability and good synchronization of the diagonal limbs (Figure 4B1, e.g., *x*_8_, *x*_9_). The variable gait of the V3^OFF^ mice (Figure 4B2) was characterized by steps with various left–right hindlimb and forelimb phase differences (e.g., see episodes *y*_1–3_ in Figure 4B2), including in-phase steps, and a high step-to-step variability.

To further understand how V3 silencing affected speed-dependent gait expression across animals, we calculated the occurrence, persistence, and attractiveness of each gait (Lemieux et al., 2016) at different speeds in WT and V3^OFF^ mice (Figure 5, Figure 5—figure supplement 1). The occurrence was the percentage of a certain gait within the total steps; the persistence of a certain gait was the likelihood of a step of this gait to be followed by another step of the same gait; the attractiveness was calculated as the number of steps transferring to the target gait divided by the total number of all circumstances of gait transition in two consecutive steps.

**Figure 5.**
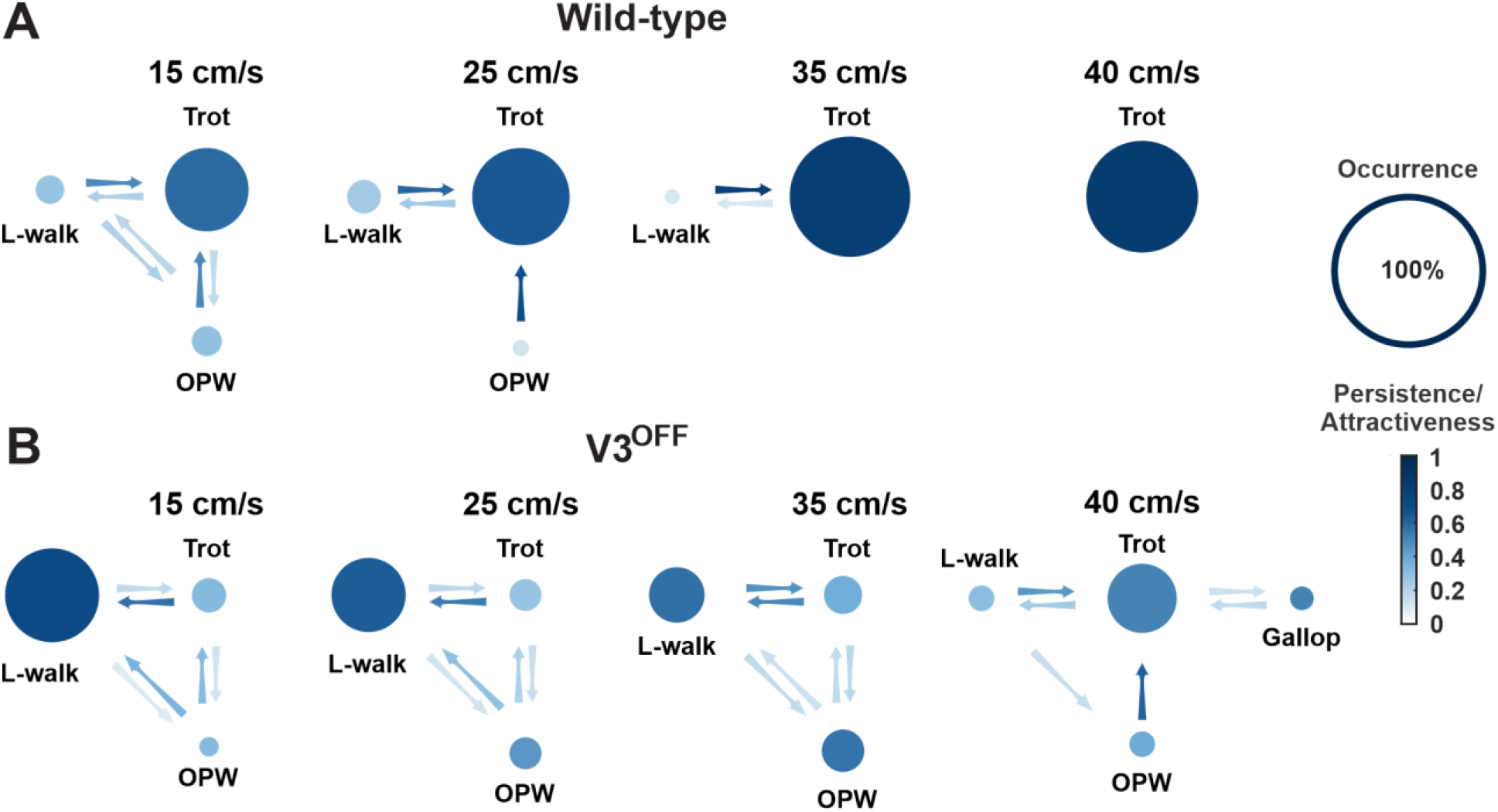
Gait transition diagrams of WT (A) and V3^OFF^ (B) at different speeds. The size of circle indicates the relative occurrence of the indicated gait. The full size of the circle meaning 100% occurrence is shown on the far right. The intensity gradient of the colour of the circles indicates the persistence of the corresponding gait. The intensity gradient of the colour of the arrows indicates the likelihood of a transition.

The relative values of these parameters for each gait and the probability of transitions between different gaits at different speeds are illustrated in Figure 5. These analyses together clearly demonstrated that gait expression of V3^OFF^ mice was altered compared to the WT mice in a speed-dependent manner (Figure 5, Figure 5—figure supplement 1). WT mice used trot as the preferred gait at all tested speeds (15–40 cm/s) and its occurrence, persistency, and attractiveness increased with speed (Figure 5A, Figure 5—figure supplement 1A). In contrast, the most prevalent, persistent, and attractive gait in V3^OFF^ mice at low to medium speeds up to 35 cm/s was a lateral-sequence walk (L-walk; Figures 5B and Figure 5—figure supplement 1B). Although V3^OFF^ animals exhibited steps that could be classified as trot, the occurrence and persistence of trot episodes was low. The difference in the preferred gait between trot in WT mice and lateral-sequence or out-of-phase walk in V3^OFF^ mice was associated with the changes in the homolateral and diagonal phase-differences (Figure 3C2, D2). As mentioned above, WT mice have trot as a dominant gait, which becomes even more prevalent and attractive with increasing speed (Figure 5A, Figure 5—figure supplement 1A). In contrast, in V3^OFF^ mice the prevalence and attractiveness of the lateral-sequence walk, representing their dominant gait, decreased with speed, while the prevalence and attractiveness of less structured out-of-phase walk increased (Figure 5B, Figure 5—figure supplement 1B). This is likely associated with the increased variability of relative phasing in V3^OFF^ mice at higher speeds.

In summary, silencing V3 neurons limited the ability of mice to locomote at high speed (Figure 2), distorted interlimb coordination, and changed homolateral and diagonal phase relationships between fore and hind limbs at low and medium speeds (Figure 3). This resulted in converting gaits in V3^OFF^ mice from trot to lateral-sequence walk at lower speeds and significantly reduced gait stability when speed increased (Figures 4 and 5).

### Modeling spinal locomotor circuits incorporating V3 neurons with local and long propriospinal projection

Our experimental results clearly indicate that V3 neurons should play important roles in speed-dependent interlimb coordination. Yet, the specific function and connectivity of different V3 subpopulations (such as V3 aLPNs and local V3 CINs) remain unknown and cannot be explicitly drawn from the current experimental data. Therefore, we extended our previous computational model of central control of interlimb coordination (Danner et al., 2017) and used it to study potential mechanisms by which the different V3 subpopulations interact with the rhythm-generating spinal circuitry and affect limb coordination.

The present computational model of spinal circuits was built based on our previous models (Danner et al., 2017, 2019). The spinal circuitry was modelled as interacting populations of neurons described in the Hodgkin-Huxley style (see Modeling methods). The rhythm-generating (RG) populations contained 200 neurons each; all other populations contained 100 neurons. Heterogeneity within the populations was ensured by randomizing the value for leakage reversal potential, initial conditions for the membrane potential, and channel kinetics variables among neurons. Interactions between and within populations were modelled as sparse random synaptic connections. Model equations, neuronal parameters, connection weights, and their probabilities are specified in Modeling methods.

The main objectives in this study were (1) to incorporate subpopulations of V3 neurons with ascending long propriospinal projections (aLPNs) from lumbar to cervical locomotor circuits, based on our tracing and *in vitro* studies, and (2) to update the model so that it could reproduce multiple effects of V3 neuron removal on limb coordination, locomotor speed, and speed-dependent gait expression observed in our experimental studies.

Similar to the previous models (Danner et al., 2016, 2017; Ausborn et al., 2019, 2021), our model has cervical and lumbar compartments, controlling fore and hind limbs, respectively (Figure 6A). Each compartment includes the left and right rhythm generators (RGs), connected by several pathways mediated by local commissural neurons (CINs). The cervical and lumbar compartments interact via homolateral and diagonal long propriospinal pathways mediated by aLPNs and dLPNs. The present model includes several distinct subpopulations of V3 neurons (Figure 6B). The local V3 CINs are involved in excitatory interactions between left and right extensor (V3-E) and left and right flexor (V3-F) half-centers within both cervical and lumbar compartments (Rybak et al., 2013, 2015; Shevtsova et al., 2015; Danner et al., 2016, 2017, 2019). The V3 aLPNs (aV3) mediate the diagonal (commissural) pathways from lumbar to cervical RGs, which have been incorporated based on our experimental results (Figure 1A1-A4).

**Figure 6.**
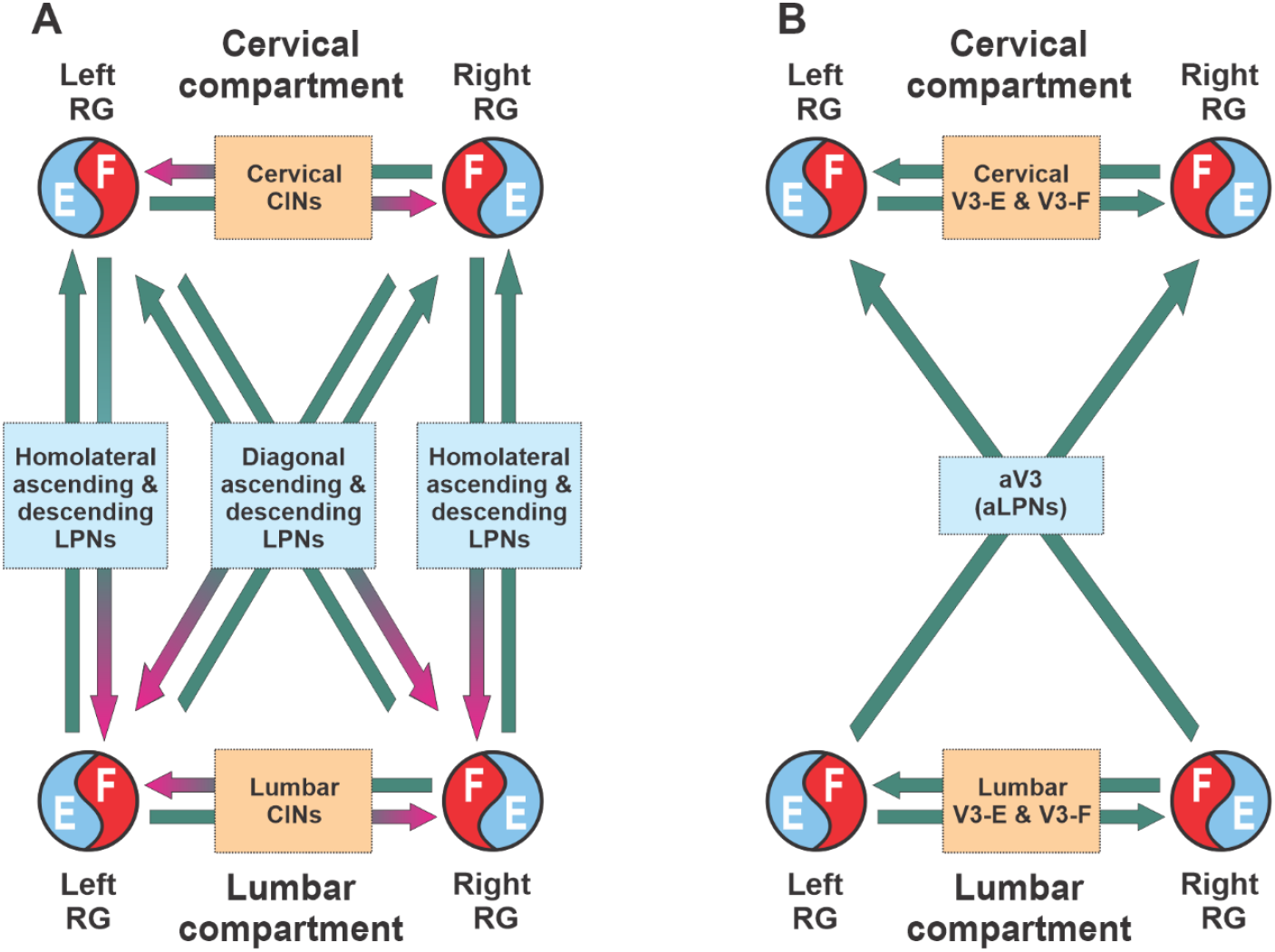
Conceptual model schematic. **A.** Model incorporates two bilateral compartments (cervical and lumbar). Each compartment includes the left and right rhythm generators (RGs) controlling each a single limb and interacting within compartment by commissural interneurons (CINs). The cervical and lumbar compartments interact via homolaterally and diagonally projecting, descending and ascending long propriospinal neurons (LPNs). **B.** Interaction between four RGs by different populations of V3 interneurons.

#### Modeling the organization of spinal circuits involved in control of locomotion

The basic architecture of the model has been developed for several years to be able to reproduce experimental data from multiple independent experimental studies (Rybak et al., 2013, 2015; Molkov et al., 2015; Shevtsova et al., 2015; Danner et al., 2016, 2017, 2019; Shevtsova and Rybak, 2016; Ausborn et al., 2019, 2021; Latash et al., 2020) and was extended here to incorporate the results of experimental studies described above. The detailed schematic of the model is shown in Figure 7 and its major components are shown in Figure 8 and described below.

**Figure 7.**
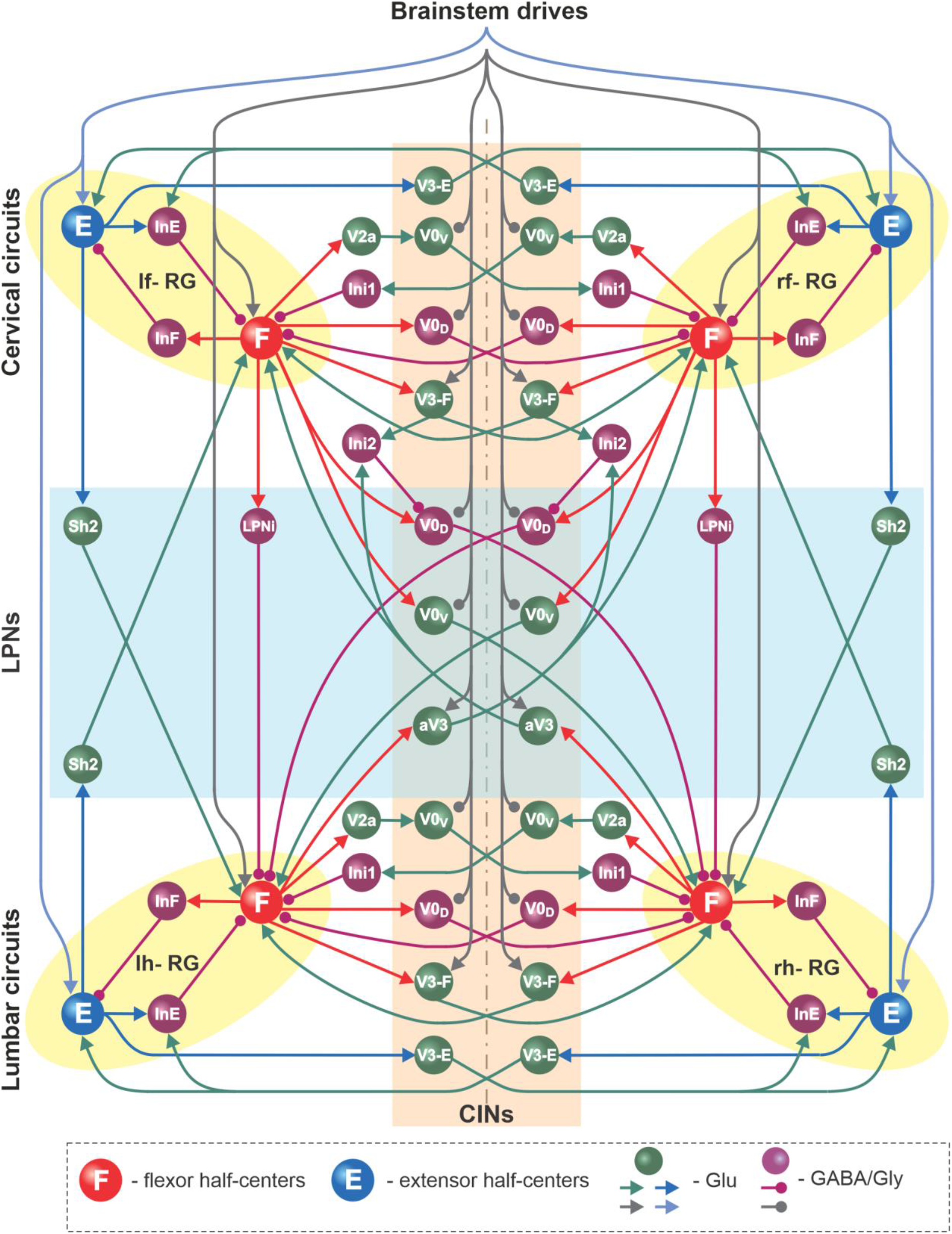
Full model schematic. Spheres represent neural populations and lines represent synaptic connections. Excitatory (Glu) and inhibitory (GABA/Gly) connections are marked by arrowheads and circles, respectively. RG, rhythm generator; CINs, commissural interneurons; LPNs, long propriospinal neurons.

**Figure 8.**
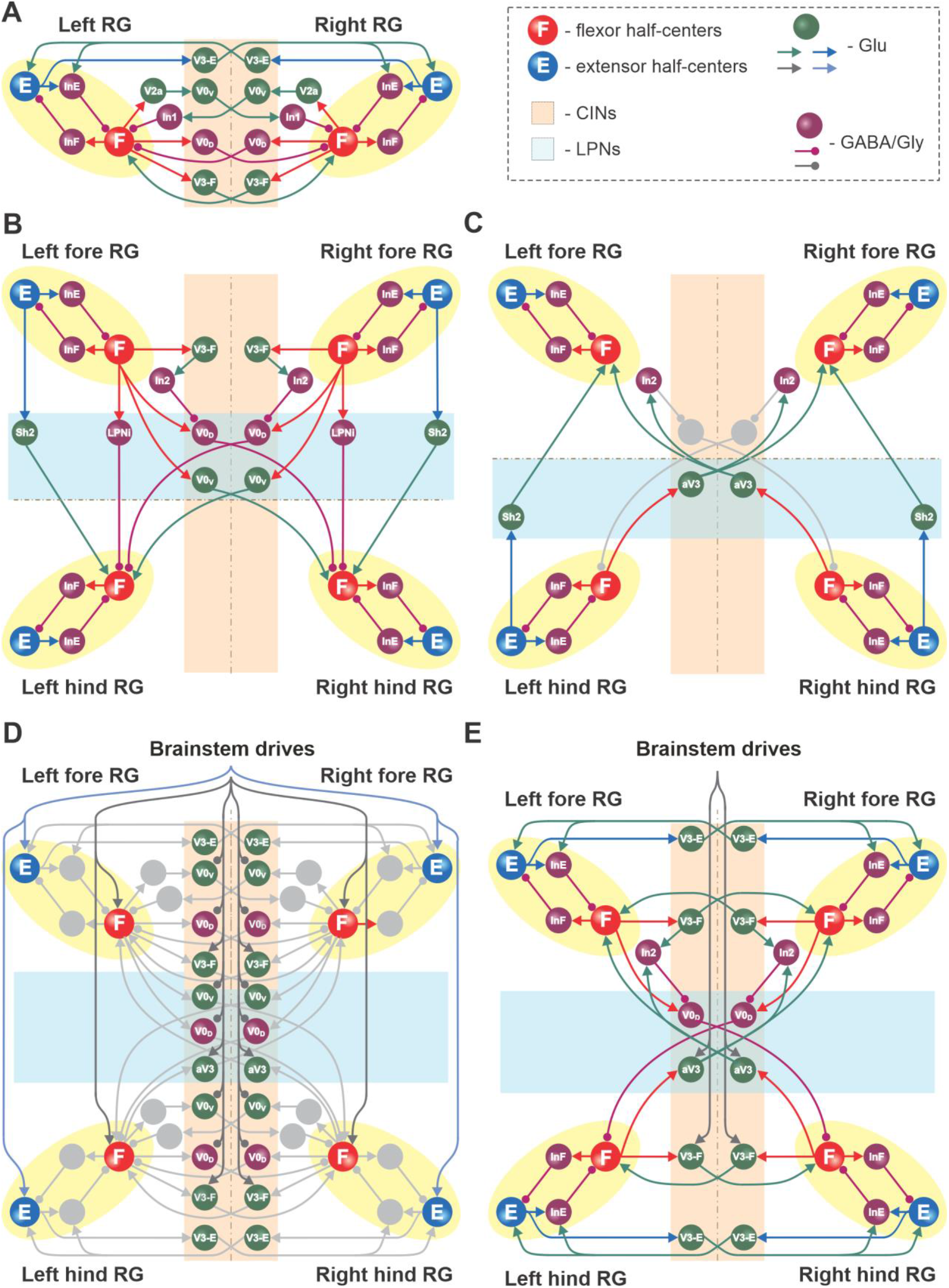
Connections within the spinal cord in the model. **A.** Connections between the left and right rhythm generators (RG) within each compartment (see text for details). **B.** Cervical-to-lumbar connections via descending long propriospinal neurons (dLPNs). **C.** Lumbar-to-cervical connections via ascending long propriospinal neurons (aLPNs). **D.** Brainstem drive to the extensor and flexor half-centers (F and E), commissural interneurons (CINs), and long propriospinal neurons (LPNs). **E.** Local and long propriospinal connections in the model mediated by V3 interneurons. Neural populations are shown by spheres. Excitatory and inhibitory connections between populations are shown by arrowheads and circles, respectively.

#### Rhythm generators (RGs)

Each of the four RGs includes the flexor (F) and extensor (E) half-centers, mutually inhibiting each other via inhibitory interneuron populations (InF and InE; Figures 7 and 8). The neurons in the F and E half-centers incorporate a slowly inactivating persistent sodium current (*I*_NaP_) and are connected by excitatory synaptic connections that allow them to generate synchronized populational bursting activity in a certain range of an external brainstem drive. As in our previous models (Rybak et al., 2013, 2015; Molkov et al., 2015; Shevtsova et al., 2015; Danner et al., 2016, 2017, 2019; Shevtsova and Rybak, 2016; Ausborn et al., 2019, 2021; Latash et al., 2020), only the F half-centers operate in a bursting regime and generate intrinsically rhythmic activity, while the E half-centers receive a relatively high tonic drive and are tonically active if uncoupled. The E half-centers generate rhythmic activity only due to inhibition from the intrinsically oscillating F half-centers. Thus, each RG generates the locomotor-like (flexor-extensor alternating) activity in a certain range of frequencies, depending on the brainstem drive to the F half-center.

#### Local commissural (left–right) interactions

Left–right interactions within cervical and lumbar compartments include several commissural pathways (Figures 8A). Two pathways are mediated by V0 (V0_V_ and V0_D_) CINs and support left–right alternating activity and alternating gaits (i.e., walk and trot). The inhibitory V0_D_ CINs provide direct mutual inhibition between the flexor half-centers. The excitatory V0_V_ CINs also provide mutual inhibition between the flexor half-centers (receiving inputs from excitatory V2a and acting through inhibitory Ini1 populations). Two other pathways mediated by two types of local excitatory V3 CINs (V3-E and V3-F) support synchronization of the left and right RG activities and promote left-right (quasi-) synchronized gaits (gallop and bound) at higher locomotor frequencies. The V3-F subpopulations provide mutual excitation between the F half-centers, similar to our previous models (Rybak et al., 2015; Shevtsova et al., 2015; Danner et al., 2016, 2017), while the V3-E CINs mediate mutual excitation between the E half-centers and are incorporated to fit our previous experimental and modeling results (Danner et al., 2019).

#### Long propriospinal interactions between cervical and lumbar circuits

The cervical and lumbar compartments interact via descending (cervical-to-lumbar) (Figure 8B) and ascending (lumbar-to-cervical) LPNs (Figure 8C) whose organization is based on our previous computational models (Danner et al., 2017; Ausborn et al., 2019, 2021). The lumbar-to-cervical pathways also incorporate ascending diagonal V3 LPNs (aV3), which are based on the present experimental data (Figure 1A1-A4).

The descending homolateral connections (Figure 8B) are mediated by the excitatory Shox2 populations (Sh2), providing excitation from each cervical E half-center to its homolateral lumbar F half-center, and by the inhibitory LPN populations (LPNi), mediating inhibition of each lumbar F half-center from the homolateral cervical F half-center. The descending diagonal connections are mediated by V0_V_ LPNs, providing excitation of each lumbar F half-center from the diagonal cervical F half-center, and by the inhibitory V0_D_ LPNs, providing inhibition of each lumbar F half-center from the diagonal cervical F half-center. The latter pathways are also regulated by the cervical V3-F populations, which inhibit the V0_D_ LPNs ipsilaterally through the inhibitory In2 populations.

The ascending connections in the model (Figure 8C) include the homolateral excitatory pathway mediated by Shox2 populations (Sh2), providing excitation from each lumbar E half-center to the homolateral cervical F half-center, and the diagonal excitatory pathway mediated by the diagonal aV3 populations, providing excitation of each cervical F half-center from the diagonal lumbar F half-center. The diagonal aV3 populations also excite the In2 populations, thus regulating activity of the diagonal V0_D_ LPNs.

#### Supra-spinal (brainstem) drive to cervical and lumbar circuits

The tonic brainstem drives are organized to excite all E and F half-centers, the local V3-F in both compartments, and the diagonal aV3 populations, and to inhibit the local V0 CINs in both compartments and the descending diagonal V0_V_ and V0_D_ LPNs (Figure 8D). The excitatory drive to all E half-centers was kept constant, while the value of the drive to the F half-centers, local CINs, and diagonal LPNs varied to provide control of locomotor frequency and the frequency-dependent gait expression (for details see Danner et al., 2016, 2017).

#### Circuit interactions mediated by V3 subpopulations

Figure 8E shows connectivity of all V3 subpopulations. In our previous models, we suggested that the local V3 CINs promote left– right synchronization by providing mutual excitation between left and right flexor (Danner et al., 2016, 2017) and left and right extensor half-centers (Danner et al., 2019; Ausborn et al., 2021). In the present model, this function is performed by V3-F and V3-E populations, respectively. Both these populations support synchronization of the left and right RG activities and promote gallop and bound at higher locomotor frequencies. Furthermore, the diagonal ascending propriospinal V3 subpopulations (aV3) mediate excitation of each cervical F half-center from the contralateral lumbar F half-center and support diagonal synchronization necessary for trot at medium locomotor frequencies. In addition, both the cervical V3-F subpopulations and the contralateral aV3 subpopulations inhibit the diagonal V0_D_ LPNs through the inhibitory In2 populations, hence securing the stable transition from walk to trot (Danner et al., 2017).

#### Model operation in the intact case

To show that the model with the updated V3 connectivity can reproduce speed-dependent gait expression of WT mice, we investigated the model behavior with all populations and connections intact. Figure 9 shows the model performance for three sequentially increased values of parameter α controlling the brainstem drive to the spinal network (see Figure 7 and 8D). Increasing the brainstem drive caused an increase in frequency of locomotor oscillations and consecutive gait transitions from lateral-sequence walk (Figure 9A1) to trot (Figure 9A2) and finally to bound (Figure 9A3). Figure 9B shows a raster plot of activity of all neurons in the left ascending propriospinal aV3 population. When the brainstem drive was progressively increased (from *α* =0 to *α* =1), the oscillation frequency increased from 2 to 8 Hz. This increase of V3 population activity with the increase of locomotor frequency (reflecting locomotor speed) was consistent with our experimental results (Figure 1C5).

**Figure 9.**
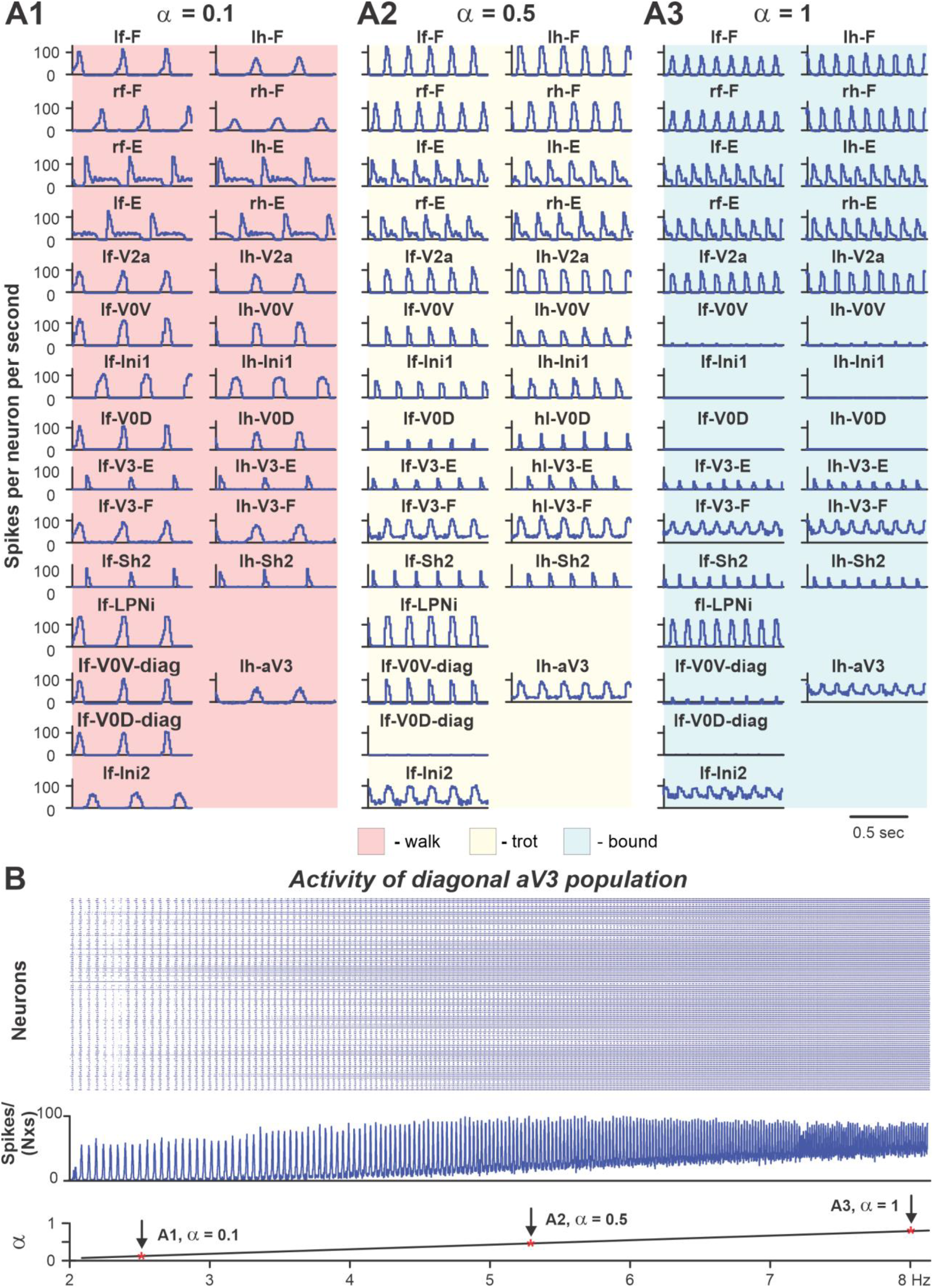
Performance of the intact model for three values of brainstem drive. In **A1–A3,** model performance for three sequentially increased values of parameter *α* (*α* = {0.1; 0.5; 1}) that controls brainstem drive to the network. In all diagrams, only left cervical (fore, f-) and lumbar (hind, h-) populations are shown. Integrated activities of all populations are shown as average histograms of neuron activity [spikes/(N × s), where N is a number of neurons in population; bin = 10 ms]. **B.** Raster plot (upper panel) of activity of the left lumbar diagonal V3 population when the brainstem drive (lower panel) was increased from 0.1 to 1. Note increased recruitment of the V3 neurons with the increased drive.

Figure 10A1 shows bifurcation diagrams reflecting changes in the normalized phase differences between oscillations in the left and right hind (lumbar) RGs, left and right fore (cervical) RGs, homolateral, and diagonal RGs with a progressive increase in brainstem drive. Increasing brainstem drive in the intact model resulted in sequential changes of gait from lateral-sequence walk to trot and then to gallop and bound (Figure 10A1) and was accompanied by increased frequency of stable locomotor oscillations (Figure 10B1). The sequential gait changes in the model were consistent with the previous experimental observations (Bellardita and Kiehn, 2015; Lemieux et al., 2016) and our modeling studies (Danner et al., 2016, 2017; Ausborn et al., 2019). During transitions from walk to trot and from trot to gallop and bound, the model exhibited bi- or multi-stable behaviors (Fig 10A1 and B1). In these areas, a small disturbance or noise could result in spontaneous transitions of model activity from one steady state regime to the other. While building the bifurcation diagrams in Figure 10, noise was added to all tonic drives in the model, and, as a result, in the range of the parameter *α* from 0.4 to 0.5, spontaneous transitions between walk and trot occurred; in the range of *α* between 0.7 and 0.85, the model exhibited either trot, gallop, or bound; and at *α* values larger than 0.85 both gallop and bound coexisted. These model behaviors correspond to previous experimental and modeling studies (Bellardita and Kiehn, 2015; Lemieux et al., 2016; Danner et al., 2017). Thus, the proposed model organization (including V3 CIN and aLPN populations and their connectivity) can account for the speed-dependent gait expression observed in WT mice.

**Figure 10.**
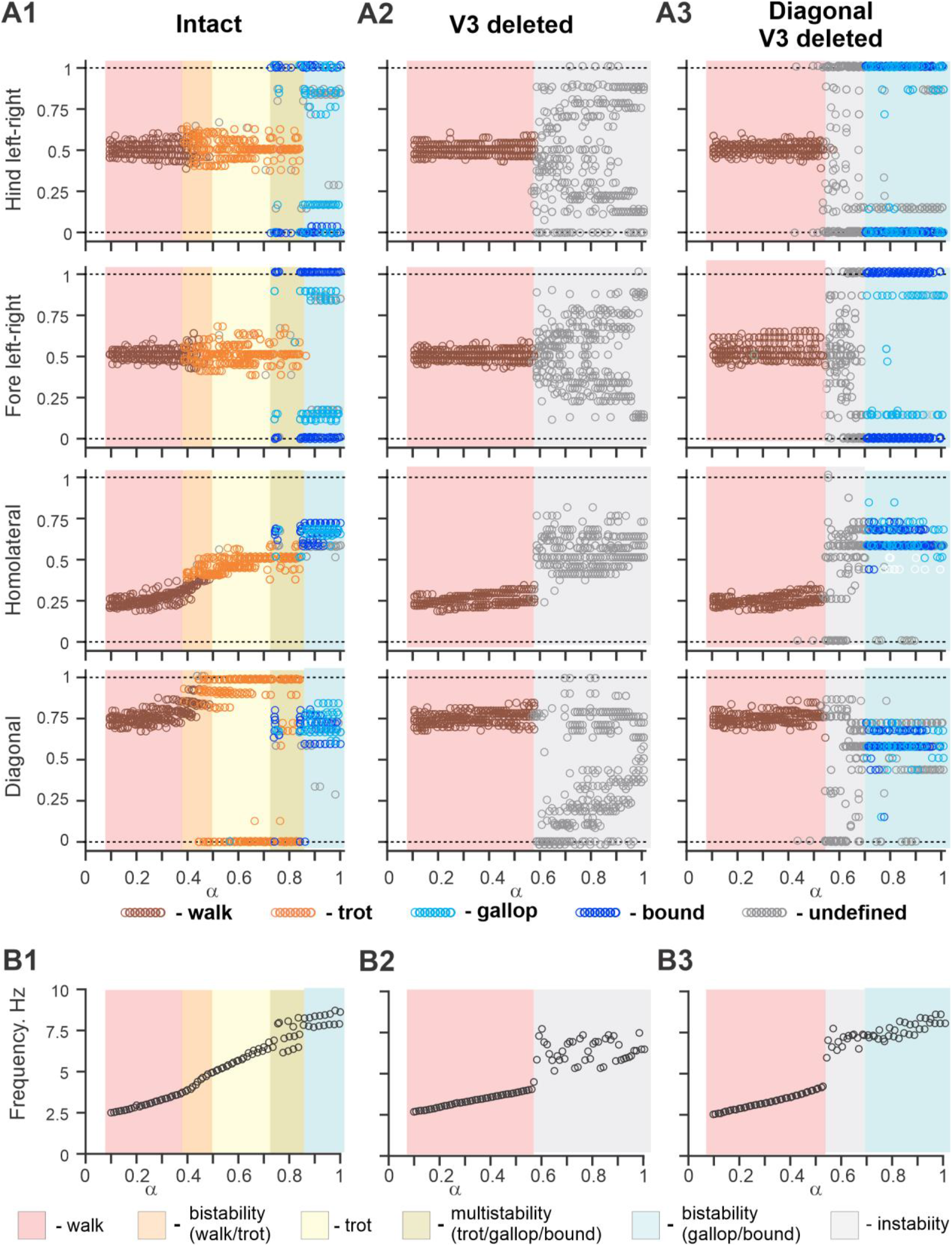
Bifurcation diagrams of the intact model, and after removal of all V3 or only propriospinal aV3 neurons. Bifurcation diagrams reflecting changes in the normalized phase differences between oscillations in the left and right hind (lumbar) RGs, left and right fore (cervical) RGs, homolateral, and diagonal RGs with a progressive increase in brainstem drive (*α*) from 0.1 to 1 with a step of 0.01 (see for details Modeling methods and Danner et al., 2017). **A1.** Intact model. **A2.** The result of removal of all V3 neurons (V3^OFF^ simulation). **A3.** Only the diagonal aV3 are removed. Parameter *α* that represents the brainstem drive was used as bifurcation parameters. Each diagram shows phase differences calculated for 5 locomotor periods when *α* was fixed. Yellow areas show ranges of *α* when the model demonstrates bi- or multistable behaviors. **B1**–**B3.** Averaged frequency corresponding to simulation shown in (**A1**–**A3**).

#### Modeling the effects of elimination of V3 neurons

The model described above was used to investigate the effects of removing the V3 population to simulate our experimental studies using V3^OFF^ mice, in which all V3 neurons were genetically silenced. Deletion of all V3 neurons produced notable changes in model operation (Figure 10A2,B2). First, the frequency range of stable locomotor oscillation was reduced, so that above the frequency of 3.8 Hz (*α*=0.57) the model transitioned to uncoordinated locomotor oscillations resulting in unstable, variable gaits (Figure 10B2). Second, the range of brainstem drive values and the range of frequencies in which the lateral-sequence walk occurred was extended, while trot and other gaits (gallop/bound) became unstable or even failed to occur (Figure 10A2). Both of these effects were qualitatively consistent with our experimental results (Figures 2 and 3).

The effects of removal V3 populations in the model are shown in Figure 11 for three values of brainstem drive defining locomotor frequency. The intact model exhibits a lateral-sequence walk at low locomotor frequencies and trot at medium frequencies (Figure 11A). After V3 removal, stable trot disappears, the model only exhibits a lateral-sequence walk at low to medium frequencies, and the interlimb coordination becomes unstable at higher frequencies, resulting in variable gaits (Figure 11B). The behavior of the intact model and the changes following the removal of all V3 neurons qualitatively corresponds to our experimental observations (Figure 4B1,B2).

**Figure 11.**
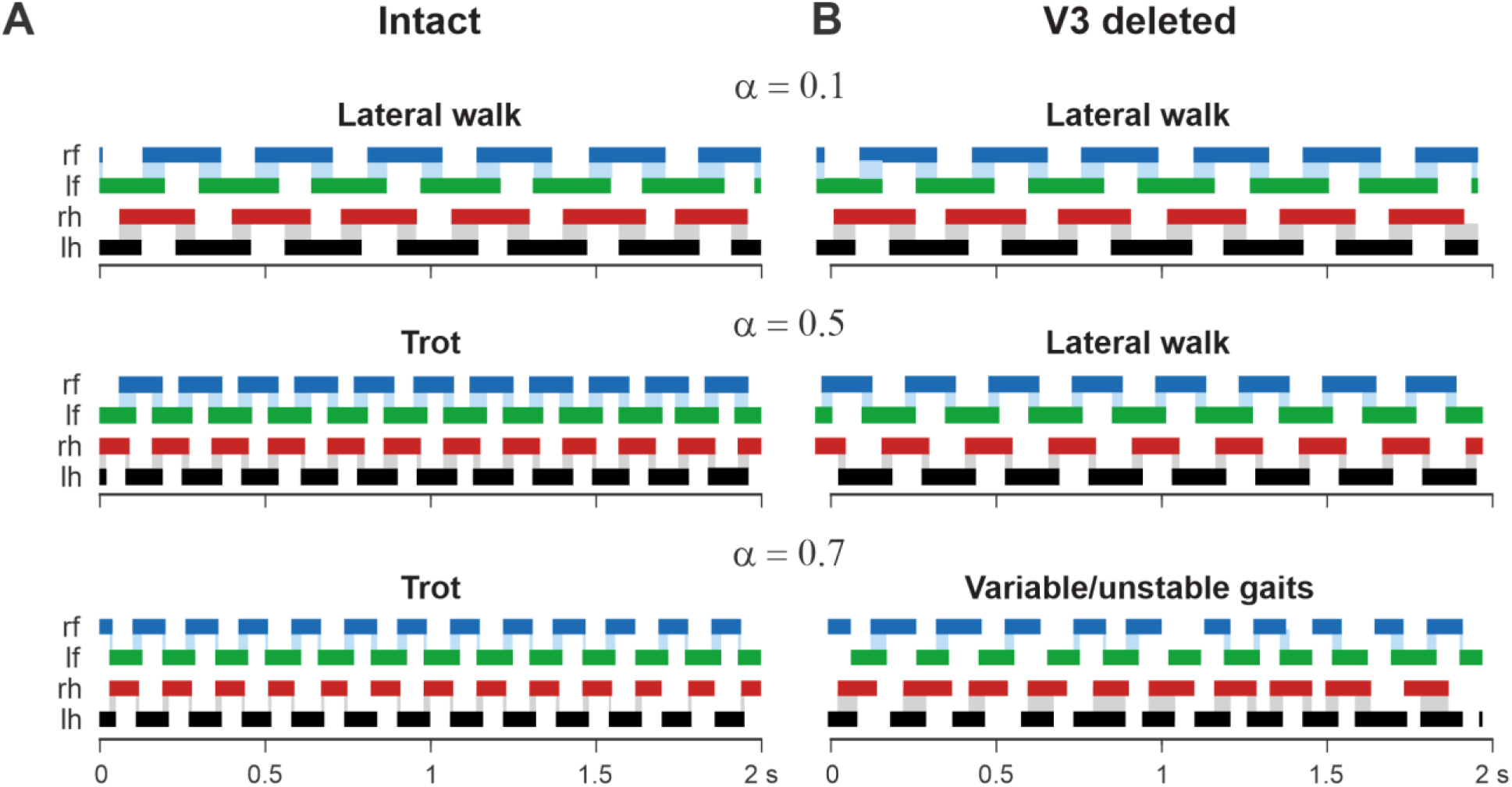
Examples of the step diagrams for the intact model (A) after all V3 neurons were deleted (B). For each rhythm generator (RG), the lengths of the corresponding extensor phases are shown for two seconds of the simulation time for three values of *α* (*α* = 0.1; *α* = 0.5; *α* = 0.7). At *α* = 0.1, both intact and V3-deleted models demonstrate stable lateral walking gait. At α = 0.5, the intact model exhibits a trot, and the V3-deleted model demonstrates a lateral-sequence walk. At *α* = 0.7, the intact model has a stable trot while the V3-deleted model demonstrates unstable behavior. Blue and gray shadows highlight periods of overlap between extensor phases of the homologue RGs.

A comparison of normalized left–right, homolateral, and diagonal phase differences in the intact model and after deletion of all V3 neurons (Figure 12) demonstrated that the intact model exhibited phase relationships typical for trot. In contrast, after removal of all V3 neurons, homolateral and diagonal phase differences were shifted towards lower values, corresponding to a lateral-sequence walk, which was similar to our experimental results (Figure 3A2-D2).

**Figure 12.**
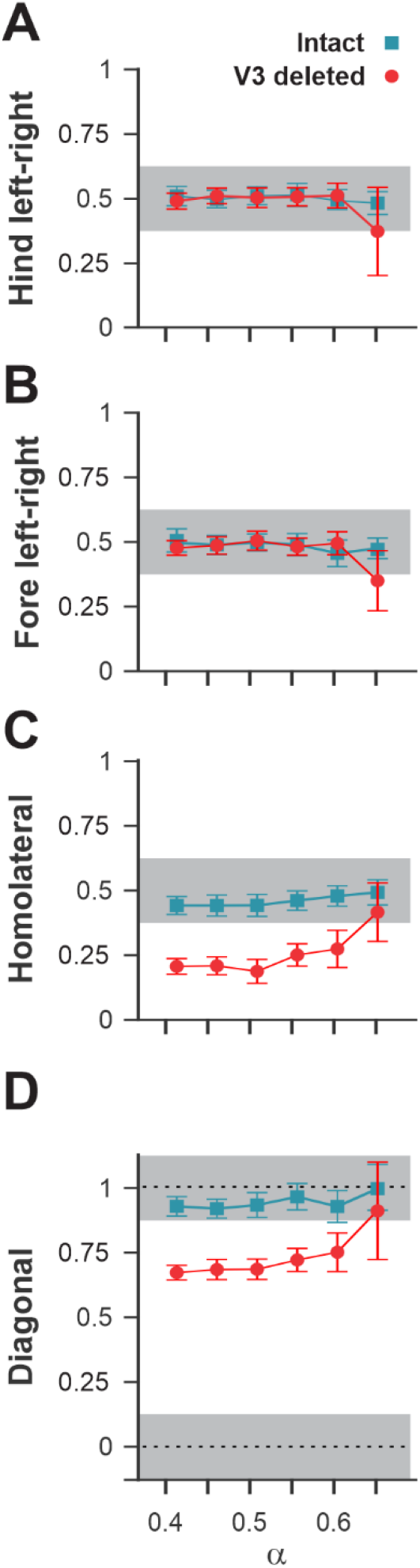
Interlimb coordination in the intact model and after removal of all V3 neurons. Interlimb coordination in the intact (blue) and V3-deleted (red) models for six values of the brainstem drive shown for left–right hind (**A**), left–right fore (**B**), homolateral (**C**), and diagonal (**D**) normalized phase differences. For each value of *α*, phase differences were averaged over 10-20 consecutive oscillation periods. In **A, C,** and **D** the extensor phase of the left hind RG is chosen as the reference. Note the homolateral and diagonal phase difference shift toward walking in V3-deleted model compared to trotting gate in the intact case.

To investigate the specific role of the ascending long propriospinal V3 (aV3) neurons, we simulated a separate case when only V3 aLPN subpopulations were removed (Figure 10 A3,B3). Our simulations have shown that the deletion of only V3 aLPNs in the model also leads to elimination of trot at medium locomotor frequencies. However, with further increases in brainstem drive, after some period of instability, sharp transitions to the (quasi-)synchronized gaits (gallop and bound) took place, which in this case were supported by the remaining local populations of V3-F and V3-E CINs. This suggests that V3 aLPNs support trot by promoting diagonal synchronization through excitation of the diagonal forelimb flexor half-centers by the hindlimb flexor half-centers (Figure 7 and 8C). This result can be considered as a modeling prediction for future experimental studies.

## Discussion

In the current study, we identified a population of ascending V3 LPNs (V3 aLPN) located in the lumbar cord that activates cervical locomotor circuits and is increasingly active with increasing locomotor speed. In mice in which the entire V3 population was silenced (V3^OFF^ mice), the maximal locomotor speed was significantly reduced, and these mice were unable to stably trot because of a lack of synchronization between diagonal pairs of limbs. Furthermore, V3^OFF^ mice exhibited high step-to-step variability in left–right coordination close to their maximum locomotor speed. We extended our previous spinal circuit model (Danner et al., 2017) to investigate the specific contribution of the individual subpopulations of V3 neurons (local CINs and aLPNs) to the speed-dependent control of interlimb coordination and gait expression. The proposed V3 aLPN connections support diagonal synchronization necessary for trot whereas the local V3 CIN connections support left–right synchronization necessary for gallop and bound. This model is able to reproduce the experimentally observed speed-dependent gait expression of WT mice as well as changes in interlimb coordination following V3 silencing. The model strongly suggests the crucial and unique roles of V3 aLPNs in control of interlimb coordination during locomotion. Finally, the model provides a potentially testable prediction for the removal of only the V3 aLPNs subset, which broadens our understanding of the locomotor circuits and will guide our future experiments.

### Lumbar V3 neurons with ascending long propriospinal projections to the contralateral cervical region

The proper coordination between limb movements during locomotion, expressed as locomotor gaits, and adaptive gait changes are essential for animals to maintain stable movement within a wide range of locomotor speeds (Hildebrand, 1976, 1980, 1989). Each limb is primarily controlled by separate rhythm-generating (RG) circuits located in a particular spinal cord compartment (Forssberg et al., 1980; Frigon, 2017). The central limb coordination is provided by multiple populations of spinal neurons that mediate mutual interactions between RG circuits controlling different limbs (Stein, 1976; Schöner et al., 1990; Danner et al., 2016, 2017; Frigon, 2017). These neuronal populations include the local commissural interneurons (CINs), operating within the cervical and lumbar enlargements and coordinating movements of left and right forelimbs and left and right hindlimbs, respectively, as well as the long propriospinal neurons (LPNs) coordinating movements of forelimbs and hindlimbs.

Our present study focused on V3 neurons, which are excitatory neurons predominately projecting to the contralateral side of the spinal cord (Zhang et al., 2008). Until now, these neurons have been primarily considered within lumbar spinal circuits. In our previous experimental and modeling studies, we suggested that V3 neurons mediate mutual excitation between left and right RG circuits and promote the left–right synchronization needed for gallop and bound (Rybak et al., 2013, 2015; Shevtsova et al., 2015; Danner et al., 2016, 2017, 2019). However, since lumbar V3 neurons diverge into several subpopulations distinguished by their distribution, morphology, and electrophysiological properties (Borowska et al., 2013, 2015; Blacklaws et al., 2015), it is likely that different V3 subpopulations perform different functions.

Here, we revealed a subpopulation of lumbar V3 neurons with ascending projections to the contralateral cervical region (V3 aLPNs). The activity of these V3 aLPNs increased with an increase of locomotor speed, suggesting their engagement during locomotion and an increase of their involvement with an increase in locomotor speed. Our model reproduced this recruitment of V3 aLPNs. Based on our modeling results, we suggest that these V3 aLPNs mediate excitation from each lumbar flexor half-center to their contralateral cervical flexor half-center, which promotes diagonal synchronization necessary for trot.

### The V3 neurons are necessary for high-speed locomotion, and their removal significantly limits the locomotor speed due to distortion of limb coordination

In our study, WT mice were able to locomote on the treadmill with high speeds up to 68–83 cm/s. Silencing V3 neurons led to a significant reduction of maximal locomotor speed. Our analysis of V3^OFF^ mouse locomotion has shown a lack of diagonal synchronization at medium speeds and an increase of variability of left–right coordination with increasing locomotor speed. Similarly, removal of all V3 neurons from our model (simulating V3^OFF^ mice) reduced the maximal frequency of stable locomotor oscillation. Model analysis revealed that this frequency reduction occurred due to the distortion of coordination between left–right homologous RGs and between diagonal lumbar and cervical RGs, both of which were mediated by V3 neurons. Yet, selective deletion of only V3 aLPNs in the model allowed for stable coordination of limb activities at high speeds, i.e., during gallop and bound, whereas trot was completely lost and the model transitioned from walk directly to gallop and bound. Together our experimental and modeling results suggest that the significant reduction of locomotor speed in V3^OFF^ mice results from the distortion of interlimb coordination, particularly the properly synchronized couplings, that are necessary for trot, gallop and bound—gaits that are normally used by WT mice at these speeds.

### The V3 neurons are involved in speed-dependent gait changes and are necessary for the expression of trot, gallop, and bound

A precise yet flexible control of interlimb coordination in a wide range of speeds allows animals to maintain dynamic stability in a continuously changing environment. This interlimb coordination, expressed as gaits, depends on, and changes with, locomotor speed and is controlled by the locomotor circuits in the spinal cord (Kiehn, 2016). Mice sequentially change gait from walk to trot and then to gallop and bound as locomotor speed increases (Clarke and Still, 1999; Herbin et al., 2004, 2007; Batka et al., 2014; Bellardita and Kiehn, 2015; Lemieux et al., 2016). WT mice use walking gaits only during low-speed exploratory locomotion and generally prefer trot for over-ground and treadmill locomotion (Gruntman et al., 2007; Bellardita and Kiehn, 2015; Lemieux et al., 2016; Caggiano et al., 2018). Gallop and bound are mainly used for escape behavior and are expressed only at high speeds (i.e., >75 cm/s), both over-ground and on the treadmill (Bellardita and Kiehn, 2015; Lemieux et al., 2016).

Our WT mice showed similar preference of the trot gait at all locomotor speeds tested (15–40 cm/s). The maximal speed investigated is below the minimal speed at which mice gallop or bound and thus these gaits were not expected to occur. In contrast to the WT mice, the V3^OFF^ mice preferred a lateral-sequence walk at almost all treadmill speeds, at which they could maintain stable limb coordination (up to 30 cm/s). At treadmill speeds close to their maximal speed (35–40 cm/s) few individual steps that could be classified as trot occurred. At these higher speeds, left–right coordination became variable and even transient episodes with left–right synchronized steps were present (Figure 4B2). Similar gait changes in left–right coordination were observed after silencing dLPNs (Ruder et al., 2016) and aLPNs of unspecific genetic type (Pocratsky et al., 2020). Our previous computational model (Danner et al., 2017) attributed the speed-dependent disruption of left–right coordination after dLPN silencing (Ruder et al., 2016) to excitatory, diagonally projecting LPNs.

In our model, the V3 aLPN populations are crucial in mediating diagonal synchronization, which is essential for trot. The local V3 CINs mediated left–right synchronization necessary for gallop and bound. The present model retained the connectivities of several other neuron types (including V0_V_, V0_D_, V2a and Shox2 neurons) from the previous model and reproduced speed-dependent gait expression (from walk to trot and then to gallop and bound) of WT mice. Elimination of all V3 populations in the model, performed to simulate locomotion of V3^OFF^ mice, extended the frequency range of the lateral-sequence walk at which the model was still able to demonstrate stable locomotor activity, eliminated trot as well as high-speed synchronous gaits, and caused variable interlimb coordination at higher speeds. Moreover, like our experimental results, the conversion of trot to lateral-sequence walk at medium locomotor frequencies resulted from a shift of both the homolateral and the diagonal phase differences. These simulation results were fully consistent with our experimental finding described above.

In summary, our experimental and modeling results allow the suggestion that the V3 neurons critically contribute to speed-dependent gait expression and are necessary for the expression of trot, gallop, and bound.

### Modeling predictions and limitations of the model

The current model includes two types of V3 neurons: local V3 CINs that operate within lumbar or within cervical compartments providing mutual excitation between the homonymous left and right RGs, and V3 aLPNs mediating diagonal excitation from each lumbar flexor half-center to the contralateral cervical flexor half-center. The local V3 CINs promote left–right synchronization necessary for gallop and bound, and the V3 aLPNs promote diagonal synchronization necessary for trot. Therefore, we suggest that the spinal cord contains two V3 subpopulations that have distinct functions. Specifically, the V3 aLPNs are necessary for the expression of trot at intermediate locomotor speeds, whereas the local V3 CINs are critical for gallop and bound at high locomotor speeds. This suggestion could be tested by silencing only V3 aLPNs: simulations predicted that this would result in a loss of trot only; gallop and bound would remain stable at higher locomotor speeds.

The other interesting feature of the model is the presence of the diagonal descending long propriospinal interaction (dLPN) mediated by the inhibitory V0_D_ neurons (Figures 7 and 8B). These connections in the model perform the opposite function to the V3 aLPNs: V0_D_ dLPNs promote diagonal alternation between the cervical and lumbar RGs and hence suppress trot and promote walk at low locomotor frequencies (low locomotor speeds). The removal of these V0_D_ neurons would make the lateral-sequence walk at low speeds unstable, which is partly supported by the experimental data *in vitro* (Talpalar et al., 2013; Kiehn, 2016). The potential roles of V0_D_ dLPN are beyond the scope of the present study. However, interestingly, to provide the stable transition from walk to trot (at slow to medium-speed locomotion), the model suggests the existence of inhibition of V0_D_ dLPNs by V3 aLPNs through intermediate inhibitory populations (In2; Figures 7 and 8C). We consider these inhibitory interactions as another modeling prediction awaiting future experimental testing.

Considering the complexity of the proposed model and current insufficiency of experimental data, the proposed model has multiple limitations. Here, we focused only on central interactions within the spinal cord, without considering biomechanics and the role of sensory feedback from the limbs which play a role in limb coordination and gait expression and stabilization. Indeed, V3 neurons are likely involved in the integration of afferent feedback. Most V3 aLPNs are located across the deep dorsal horn and intermediate region of the rostral lumbar spinal cord. Our previous study showed that V3 neurons in these regions are highly active during overground activity, but silent during swimming (Borowska et al., 2013). Neurons in this region are also known to receive intensive innervation from group Ib, group II and skin sensory afferents (Edgley and Jankowska, 1987; Bannatyne et al., 2009; Jankowska and Edgley, 2010). Therefore, the lumbar V3 aLPNs may be involved in mediating and integrating signals from both the lumbar locomotor RGs and sensory information from hindlimb muscles and joints to regulate the coordination of forelimb and hindlimb movements during locomotion. Future studies may consider afferent feedback interactions with V3 neurons and other parts of the spinal locomotor circuitry by integrating the neural network models with a model of the musculoskeletal system (Nishikawa et al., 2007; Markin et al., 2016; Prilutsky and Edwards, 2016; Ausborn et al., 2021) to simulate interactions between the neural circuits, biomechanical constrains, and the environment. We also did not consider spinal circuits operating below the RGs, such as pattern formation networks, circuits related to muscle afferent input, and reflex circuits including Ia, and Ib interneurons, Renshaw cells, and motoneurons (see Rossignol et al., 2006; Rybak et al., 2006a, 2006b; Alvarez and Fyffe, 2007; McCrea and Rybak, 2007, 2008; Pierrot-Deseilligny and Burke, 2012; Zhong et al., 2012), which play an important role in intralimb coordination and could potentially also affect interlimb coordination but remain for future study.

Nevertheless, here, we take advantage of available experimental and computational tools to demonstrate that two subgroups of lumbar V3 neurons, which possess local projections or long ascending propriospinal commissural projections, are essential components in the locomotor circuits that promote speed-dependent interlimb synchronization to perform trot, gallop, and bound.

## Material and Methods

### Experimental materials and methods

#### Animals

The generation and genotyping of Sim1^Cre/+^ mice were described previously by Zhang et al. (2008). Conditional knock-out of vGluT2 in Sim1-expressing V3 INs, Sim1^cre/+^; vGluT2^flox/flox^ (V3^OFF^ mice), was described previously by Chopek et al. (2018). Ai32 mice (Gt(ROSA)26^floxstopH134R/EYFP/+^, Jackson Laboratory, Stock No. 012569) were crossed with Sim1^Cre/+^ to generate Sim1^Cre/+^; Ai32 mice. TdTomato Ai9 mice (Rosa26^floxstopTdTom^, Jackson Laboratory, Stock No. 007909) were crossed with Sim1^Cre/+^ to generate Sim1^Cre/+^; Rosa26^floxstopTdTom^ (Sim1TdTom) mice (Blacklaws et al., 2015). All procedures were performed in accordance with the Canadian Council on Animal Care and approved by the University Committee on Laboratory Animals at Dalhousie University.

#### Treadmill locomotion

Treadmill locomotion tests were performed using 11 (6 males, 5 females) V3^OFF^ (Sim1^cre/+^;VGluT2^flox/flox^) mice and 7 (2 males, 5 females, Sim1^+/+^;vGluT2^flox/flox^ or Sim1^+/+^;vGluT2^+/flox^) control littermates (WT) at postnatal day 40 to 48 (P40–48). No training was performed before any locomotion tests. During the locomotion tests, the mice were subjected to a treadmill, Exer Gait XL, (Columbus Instruments), at speeds from 15 cm/s to 40 cm/s in 5 cm/s increments. To avoid their fatigue, animals would not perform more than two trials for each speed during each experiment, and each trail was <20 s. There was a >1 min rest period between consecutive trials.

A mirror was placed underneath the transparent treadmill belt with 45 degrees angle to project the image of the paws to a high-speed camera. The paw movements were captured at 200 frames/s. Episodes with at least 10 consecutive steps were included for analysis. For the maximum speed test, if the mouse failed to hold the speed more than three seconds after five tries, this speed was defined as the maximum speed for this mouse.

#### Gait analysis

The movement of four paws in the videos was manually tracked using Vicon Motus software. The foot contact was defined as the time of the first frame in three consecutive frames in which the size of the image of the paw on the ground did not change. The foot lift was defined as the time of the frame when the front part of the paw disappeared from the image. The time sequences of the paw movement were exported and processed using a custom-written script in Spike2 (Version 7.09a, Cambridge Electronic Design). The stride duration was defined as the duration between two consecutive foot contacts of the same paw. A stance phase started at the foot contact and ended at the foot-lift. A swing phase was the period from the offset of stance to the onset of the next stance. The normalized phase differences between the limbs were calculated as the time difference between stance onset of the tested limb and the reference limb (left hindlimb; lh) divided by the step-cycle duration of the reference limb (Figure 4A). All normalized phase differences ranged from 0 to 1. The value of 0 or 1 indicated a perfect in-phase coupling (synchronization), while 0.5 indicated a perfect anti-phase coupling (alternation). We classified trot, lateral-sequence walk (L-walk), bound, and out-of-phase walk (OPW) gaits as described in Lemieux et al. (2016). All other gaits defined by Lemieux et al. (2016) were categorized as ‘others’.

To describe the role of different gaits during treadmill locomotion, we calculated their occurrence, persistence, and attractiveness in all steps (Lemieux et al. 2016). The occurrence was the percentage of a given gait within the total steps of a trial. The persistence of a certain gait was the percentage of two consecutive steps using the same gait out of all the consecutive steps. The attractiveness was the possibility of other gaits transferring to the focused gait.

#### CTB injection and detection of c-Fos expression

Retrograde tracer, cholera-toxin B (CTB), was injected into C5 to C8 segments of P30–35 Sim1TdTom mice to detect the cervical projection of V3 INs in different spinal cord regions. Briefly, the mouse was anesthetized under isoflurane throughout the surgery. A 1-cm sagittal incision was made centered over the sixth cervical spinous process. A laminectomy was performed, and the bone covering the upper half of the sixth cervical segment (C6) was removed. The glass micropipette filled with the tracer at the tip was lowered into the spinal cord. The manipulator adjusted the position of the micropipette. 500–700 nl of CTB was injected to one side of the spinal cord using a Nanoject II (Drummond). The pipette was the kept in place for an additional 5 minutes for diffusion. After injection, the muscles were sewn back together in layers with absorbable sutures, and the skin was sewn closed with polypropylene surgical sutures.

After 7 days for the tracer transport, the animals were subjected to walk (15 cm/s) or run (40 cm/s) on a treadmill for 3 times 15 minutes with a 5-minute interval between trials. The animals in the control group were left in the home cage through the experiments. After the task, mice were put back to the home cage for 60 mins. Then, the animals were perfused with 4% paraformaldehyde (PFA). Spinal cords were harvested and postfixed in 4% PFA at 4°C for 3 hours and then cryoprotected in 20% sucrose before embedding in OCT and cryostat sectioning. The 30 μM thick sections were collected for immunolabeling. Sections were washed in 0.01 M PBS with 0.1% Triton X-100 (PBS-T), blocked for 1 hour in 5% normal goat serum in PBS-T, and incubated at room temperature over two nights in rabbit anti-cFos antibody (1:2000; Santa Cruz Biotechnology) in PBS-T with 2% goat serum. Sections were washed in 3 X PBS-T and incubated with goat anti-rabbit conjugated to Alexa Fluor 488 (1:500; The Jackson Laboratory) for 2 hours at room temperature. Sections were then washed in PBS three times and coverslipped with an anti-fade mounting medium (Dako).

The images were obtained using a Zeiss LSM 510 upright confocal microscope or a Zeiss Axiovert 200M fluorescent microscope. Total tdTomato-positive V3 neurons and those co-expressing CTB and Fos protein were counted and mapped. For each animal, we sampled a total of 20 sections from the lumbar region and summarized the relative position of all double-labelled CTB/tdTomato cells and triple-labelled CTB/tdTomato/c-Fos cells onto one schematic cross-section. In addition to using the central canal and Rexed's laminae as landmarks, we also set grids on the image of transverse sections of the spinal cord and the schematic section to map the double-labelled cells more accurately.

#### Electrophysiology

All experiments were performed using spinal cords from Sim1Ai32 mice at P2–P3. The mice were anesthetized, and the spinal cords caudal to C1 were dissected out in Ringer’s solution at room temperature (111 mM NaCl, 3.08 mM KCl, 11 mM glucose, 25 mM NaHCO_3_, 1.25 mM MgSO_4_, 2.52 mM CaCl_2_, and 1.18 mM KH_2_PO_4_, pH 7.4). The spinal cord was then transferred to the recording chamber to recover at room temperature for at least 1 hour before recording in Ringer’s solution. The recording chamber was partitioned by a narrow petroleum jelly (Vaseline) bridge into two parts with independent perfusion systems. The T6–T8 spinal segments were in the petroleum jelly and the whole lumbar (L) and cervical (C) region were exposed in the bath. A dye was added into one of the compartments to check the water tightness in the end of experiment. Electroneurogram (ENG) recordings of the L5 and C5/C8 ventral roots were conducted using differential AC amplifier (A-M system, model 1700) with the band-pass filter between 300 Hz and 1 kHz. Analog signals were transferred and recorded through the Digidata 1400A board (Molecular Devices) under the control of pCLAMP10.3 (Molecular Devices).

To activate ChR2 in V3 INs, 488 nm fluorescent light was delivered by Colibri.2 illumination system (Zeiss) through 10x 1.0 numerical aperture (NA) objectives mounted on an up-right microscope (Examiner, Zeiss) onto the ventral surface of L2–L3 segments of the isolated spinal cord. Continuous light stimuli with duration of 200 ms were used. The stimulation was applied 10 times. Then 2 mM kynurenic acid (Sigma-Aldrich) was added into the compartment for the lumbar spinal cord. The same optical stimulation to the same area was applied during the drug application and after the washout.

#### Statistical analysis

The statistical analysis was performed in Prism7 (GraphPad Software, Inc.) and MATLAB (Version R2018a, The MathWorks, Inc.). Kolmogorov-Smirnov test was used to compare the difference of maximum speed between WT and V3^OFF^ mice. Welch’s t-test were used to compare the difference of c-Fos expression in Sim1 cells and Sim1/CTB cells among different conditions. Watson-Williams test of Homogeneity of Means was used to compare phase differences between WT and V3^OFF^ mice across all speeds and at specific speeds. The Equal Kappa test was used to compare the concentration (a measure of variability) of the phase-differences between WT and V3^OFF^ mice across all speeds and at specific speeds. P-values of all post-hoc tests (comparisons at specific treadmill speeds) were adjusted using Bonferroni’s method.

## Modeling methods

### Single neuron

All neurons were simulated in the Hodgkin-Huxley style as single-compartment models. The membrane potential, *V* in neurons of the flexor (F) and extensor (E) half-centers was described as

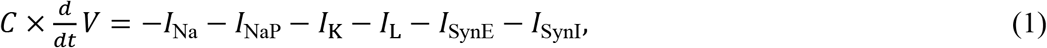

where *C* is the membrane capacitance; *t* is time; *I*_Na_ is the fast Na^+^ current; *I*_NaP_ is the persistent (slowly-inactivating) Na^+^ current; *I*_K_ is the delayed-rectifier K^+^ current; *I*_L_ is the leakage current; and *I*_SynE_ and *I*_SynI_ are synaptic excitatory and inhibitory currents.

In all other populations, the neuronal membrane potential was described as follows:

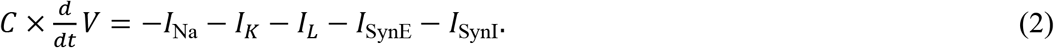

The ionic currents in Equations (1) and (2) were described as follows:

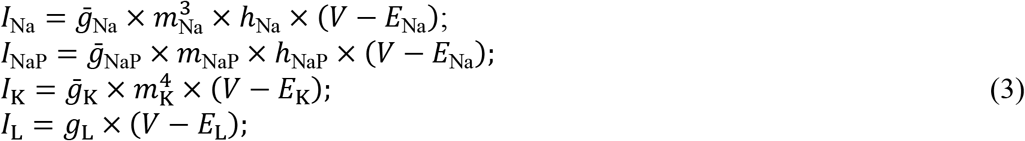

where 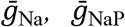 (present only in RG neurons), 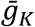, and *g*_L_. are maximal conductances of the corresponding currents; *E*_Na_, *E*_K_, and *E*_L_ are the reversal potentials for Na^+^, K^+^, and leakage currents, respectively; variables *m* and *h* are the activation and inactivation variables of the corresponding ionic channels (indicated by the indices). The maximal conductances for ionic currents and the mean leak reversal potentials, *E*_L0_, were defined as follows: in F and E half-centers, 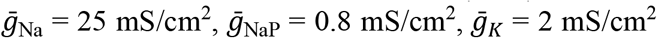 and *g*_L_ = 0.18 mS/cm^2^, *E*_L0_ = −66.4(±0.664) mV; in all other populations, 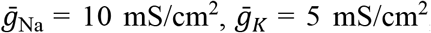, and *g*_L_ = 0.1 mS/cm^2^, *E*_L0_ = −68(±2) mV.

Activation *m* and inactivation *h* of voltage-dependent ionic channels (i.e., Na, NaP, and K) in Equation (3) were described by the differential equations:

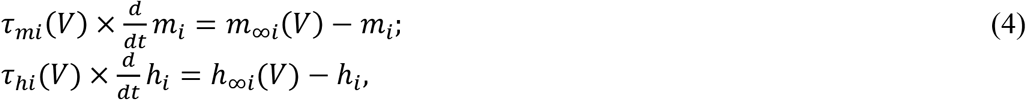

where *m*_∞*i*_(*V*) and *h*_∞*i*_(*V*) define the voltage-dependent steady-state activation and inactivation of the channel *i*, respectively, and *τ*_*mi*_(*V*) and *τ*_*hi*_(*V*) define the corresponding time constants. Activation of the sodium channels was considered instantaneous. The expressions for channel kinetics in Equation (4) are described as follows:

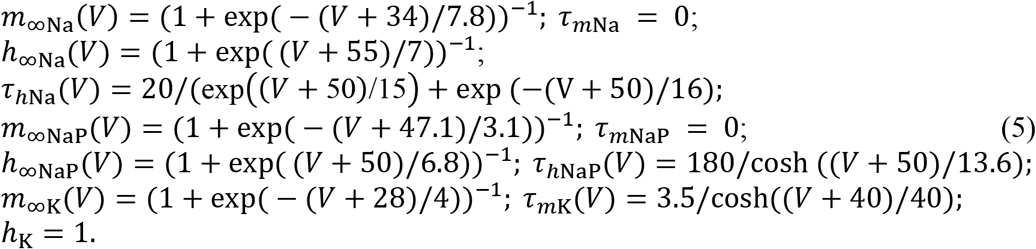

The synaptic excitatory (*I*_SynE_ with conductance *g*_SynE_ and reversal potential *E*_SynE_) and inhibitory (*I*_SynI_ with conductance *g*_SynI_ and reversal potential *E*_SynI_) currents were described as follows:

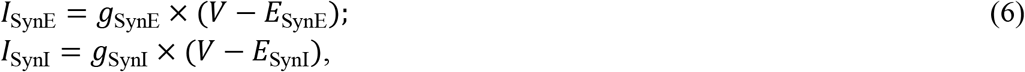

where *g*_SynE_ and *g*_SynI_ are equal to zero at rest and are activated by the excitatory or inhibitory inputs, respectively:

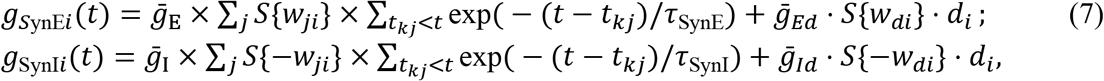

where *S*{*x*} = *x*, if *x* ≥ 0, and 0 if *x* < 0. In Equation (7), the excitatory and inhibitory synaptic conductance have two terms: one describing the effects of excitatory or inhibitory inputs from other neurons in the network and the other describing effects of inputs from the external brainstem excitatory or inhibitory drives (see also Rybak et al., 2006a). Each spike arriving to neuron *i* in a target population from neuron *j* in a source population at time *t*_*kj*_ increases the excitatory synaptic conductance by 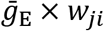 if the synaptic weight *w*_*ji*_ > 0 or increases the inhibitory synaptic conductance by 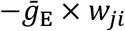 if the synaptic weight *w*_*ji*_ < 0. 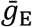 and 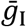 define an increase in the excitatory or inhibitory synaptic conductance, respectively, produced by one arriving spike at |*w*_*ji*_| = 1. *τ*_SynE_ and *τ*_SynI_ are the decay time constants for *g*_SynE_ and *g*_SynI_, respectively. In the second terms of equations (7), 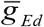 and 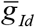 are the parameters defining the increase in the excitatory or inhibitory synaptic conductance, respectively, produced by external input drive *d*_*i*_ = 1 with a synaptic weight of |*w*_*di*_| = 1.

Excitatory and inhibitory drives to population *i* were modeled as a linear function of the free parameter *α*:

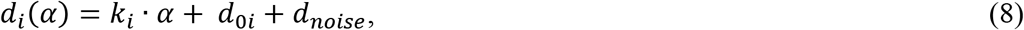

where *k*_*i*_ is the slope, *d*_0*i*_ is the intercept, and *α* is the strength of the brainstem drive. The values of *k*_*i*_ and *d*_0_ for all populations receiving the brainstem drive are indicated in Table 1. For the calculation of the bifurcation diagrams (Figure 10), Gaussian noise with a mean of 0 and a standard deviation of 10% of the drive amplitude was added to the drive to characterize the model stability and facilitate transition between gaits in the areas of bi- and multistability.

**Table 1.** Brainstem drive parameters.

| Target population | $k_i$ | $d_{0i}$ |
| --- | --- | --- |
| *Excitatory drives* |  |  |
| RG-E | 0.0 | 0.5 |
| RG-F (f) | 0.15 | 0 |
| RG-F (h) | 0.3 | 0 |
| V3-F | 1 | 0 |
| aV3 | 1 | 0 |
| *Inhibitory drives* |  |  |
| V0<sub>v</sub> | -8 | 0 |
| V0<sub>D</sub> | -10 | 0 |
| Diagonal V0<sub>v</sub> | -3 | 0 |
| Diagonal V0<sub>D</sub> | -3 | 0 |

(f) - forelimb, (h) - hindlimb, RG - rhythm generator.

The following general neuronal parameters were used: *C* =1 μF·cm^−2^; *E*_Na_ = 55 mV; *E*_K_ = − 80 mV; *E*_SynE_ = −10 mV; *E*_SynI_ = −70 mV; 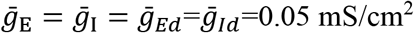; *τ*_SynE_ = *τ*_SynI_ = 5 ms.

### Neuron populations

The F and E half-centers in the model had 200 neurons, all other populations incorporated 100 neurons. Random synaptic connections between the neurons of interacting populations were assigned prior to each simulation based on probability of connection, *p*, so that, if a population *A* was assigned to receive an excitatory (or inhibitory) input from a population *B*, then each neuron in population *A* would get the corresponding synaptic input from each neuron in population *B* with the probability *p*{*A*, *B*}. If *p*{*A*, *B*}<1, a random number generator was used to define the existence of each synaptic connection; otherwise (if *p*{*A*, *B*}= 1) each neuron in population *A* received synaptic input from each neuron of population *B.* Values of synaptic weights (*w*_*ji*_) were set using random generator and were based on average values of these weights 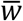 and variances, which were define as 5% of 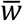 for excitatory connections 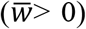 and 10% of *w* for inhibitory connections 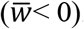. The average weights and probabilities of connections are specified in Table 2.

**Table 2.** Connection weights.

| Source | Target ($w_{ij}$) |
| --- | --- |
| Within cervical and lumbar circuits |  |
| i-F | i-F (0.0125, $p=0.1$); i-InF (0.3, $p=0.1$), i-V0<sub>D</sub> (0.3, $p=0.1$), i-V2a (0.3, $p=0.1$), i-V3-F (0.3, $p=0.1$) |
| i-E | i-E (0.0125, $p=0.1$); i-InE (0.2, $p=0.1$), i-V3-E (0.2, $p=0.05$), i-Sh2 (0.1, $p=0.1$) |
| i-IniF | i-E (-0.6, $p=0.1$) |
| i-IniE | i-F (-0.07, $p=0.1$) |
| i-V2a | i-V0<sub>V</sub> (0.4, $p=0.1$) |
| i-V0<sub>V</sub> | c-Ini1 (0.4, $p=0.1$) |
| i-Ini1 | i-F (-2, $p=0.1$) |
| i-V0<sub>D</sub> | c-F (-1, $p=0.1$) |
| i-V3-E | c-E (0.01, $p=0.05$), c-InE (0.2, $p=0.05$) |
| i-V3-F | c-F (0.03, $p=0.1$), i-Ini2 (0.05, $p=0.1$) |
| Within cervical circuits |  |
| i-F | i-LPNi (0.4, $p=0.1$), diagonal i-V0<sub>D</sub> (0.25, $p=0.1$), diagonal i-V0<sub>V</sub> (0.25, $p=0.1$) |
| Within lumbar circuits |  |
| i-F | i-aV3 (0.1, $p=0.1$) |
| Between cervical and lumbar circuits |  |
| i-Sh2 (f) | ih-F (0.02, $p=0.1$) |
| i-Sh2 (h) | if-F (0.05, $p=0.1$) |
| i-LPNi (f) | ih-F (-0.2, $p=0.1$) |
| diagonal i-V0<sub>D</sub> (f) | ch-F (-1.2, $p=0.1$) |
| diagonal if-V0<sub>V</sub> (f) | ch-F (0.1, $p=0.1$) |
| i-aV3 | cf-F (0.12, $p=0.1$), c-Ini2 (0.5, $p=0.1$) |

i - ipsilateral, c - contralateral, (f), f- forelimb, (h), h - hindlimb

Heterogeneity of neurons within each population was provided by random distributions of the mean leakage reversal potentials 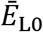 and initial conditions for the values of membrane potential and channel kinetics variables. The values of *E*_L0_ and all initial conditions were assigned prior to simulations from their defined average values and variances using a random number generator. A settling period of 1–5 s was allowed in each simulation to stabilize the model variables.

### Computer simulations

All simulations were performed using the custom neural simulation package NSM 2.5.11. The simulation package and model configuration file to create the simulations presented in the paper are available at https://github.com/RybakLab/nsm (will be uploaded after revision). The simulation package was previously used for the development of several spinal cord models (Rybak et al., 2006a, 2006b, 2013; McCrea and Rybak, 2007, 2008; Zhong et al., 2012; Shevtsova et al., 2015; Shevtsova and Rybak, 2016). Differential equations were solved using the exponential Euler integration method with a step size of 0.1 ms. Simulation results were saved as ASCII files containing time moments of spikes for all RG populations.

### Data analysis in computer simulations

The simulation results were processed using custom Matlab scripts (The Mathworks, Inc., Matlab 2020b). To assess the model behavior, the averaged integrated activities of F and E half-centers (average number of spikes per neuron per second) were used to determine the onsets and offsets of flexor and extensor bursts and calculate the flexor and extensor burst durations and oscillation period. The timing of onsets and offsets of flexor and extensor bursts was determined at a threshold level equal to 10% of the average difference between maximal and minimal burst amplitude for a particular RG population in the current simulation. The locomotor period was defined as the duration between two consecutive onsets of the extensor bursts. Duration of individual simulations depended on the value of parameter *α*, and to robustly estimate average values of burst duration and oscillation. For each value of *α*, the first 10–20 transitional cycles were omitted to allow stabilization of model variables.

Normalized phase-differences were calculated as the durations between the onsets of the extension phase of each RGs and the reference (left hind) RG divided by the period and averaged for the 10–20 consecutive cycles (see above). Gait classification was performed as for the experimental data using the definitions of Lemieux et al. (2016). To evaluate possible gaits with increasing brainstem drive, parameter *α* was linearly increased and for each value of *α* the average frequency and left–right, homolateral, and diagonal phase differences were calculated. The frequency and phase differences were then plotted against the parameter *α*.

---

**Figure S3—figure supplement 1.**
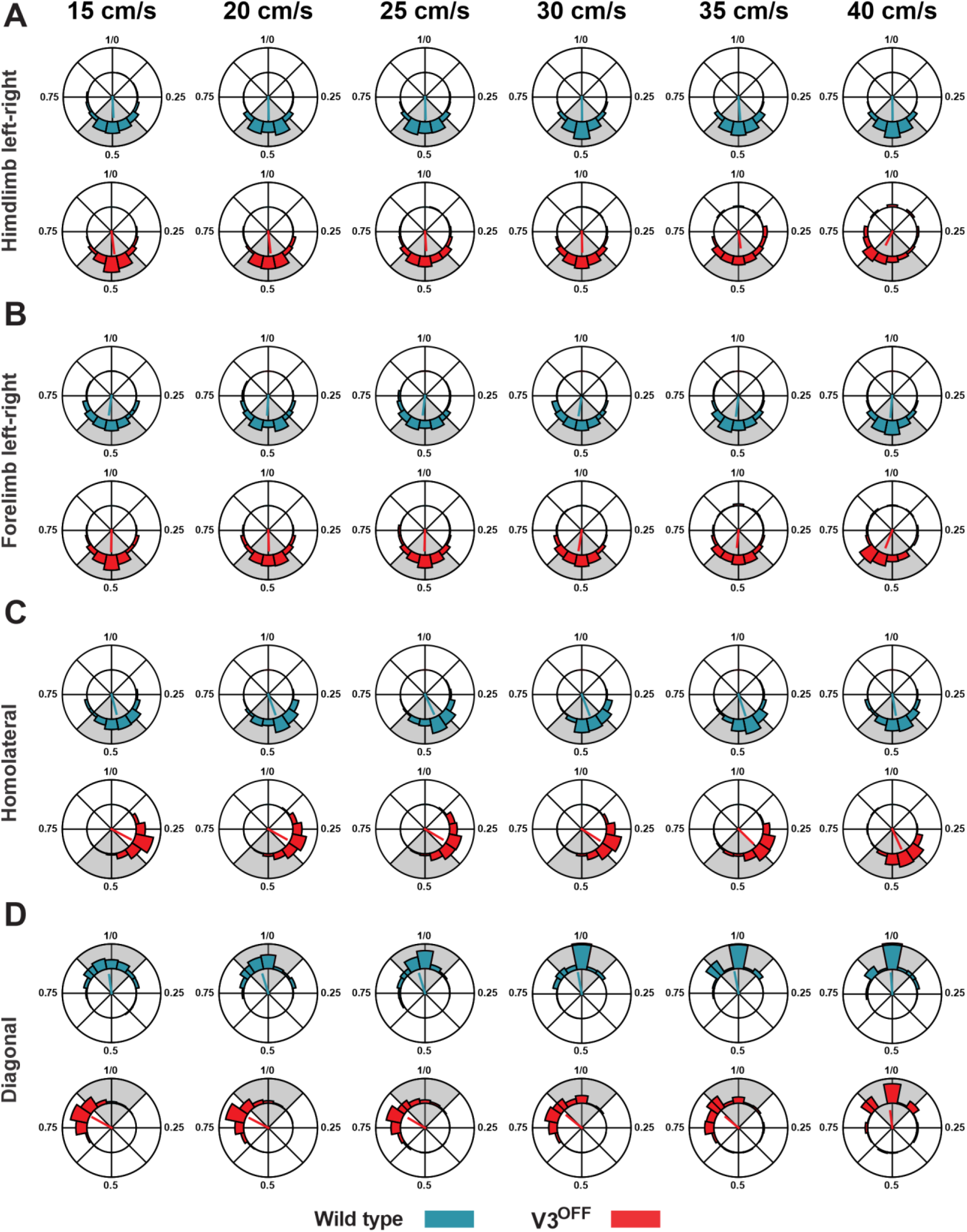
Limb couplings of WT and V3OFF mice at different speeds. Circular plots of hindlimb (**A**) and forelimb (**B**) left–right phase differences, homolateral phase differences (**C**) and diagonal phase differences (**D**) in wild type (WT; blue) and V3^OFF^ (red) mice at individual treadmill speeds. Except for the forelimb left–right phase differences, the left hindlimbs are used as the reference limb. Each vector, blue line (WT) and red line (V3^OFF^), in the circular plot, indicates the mean value (direction) and robustness (radial line/length) of the phase differences. The circle is evenly separated into 8 fractions. The circular histograms represent the distribution of phase differences of all steps at all tested speeds

**Figure S5—figure supplement 1.**
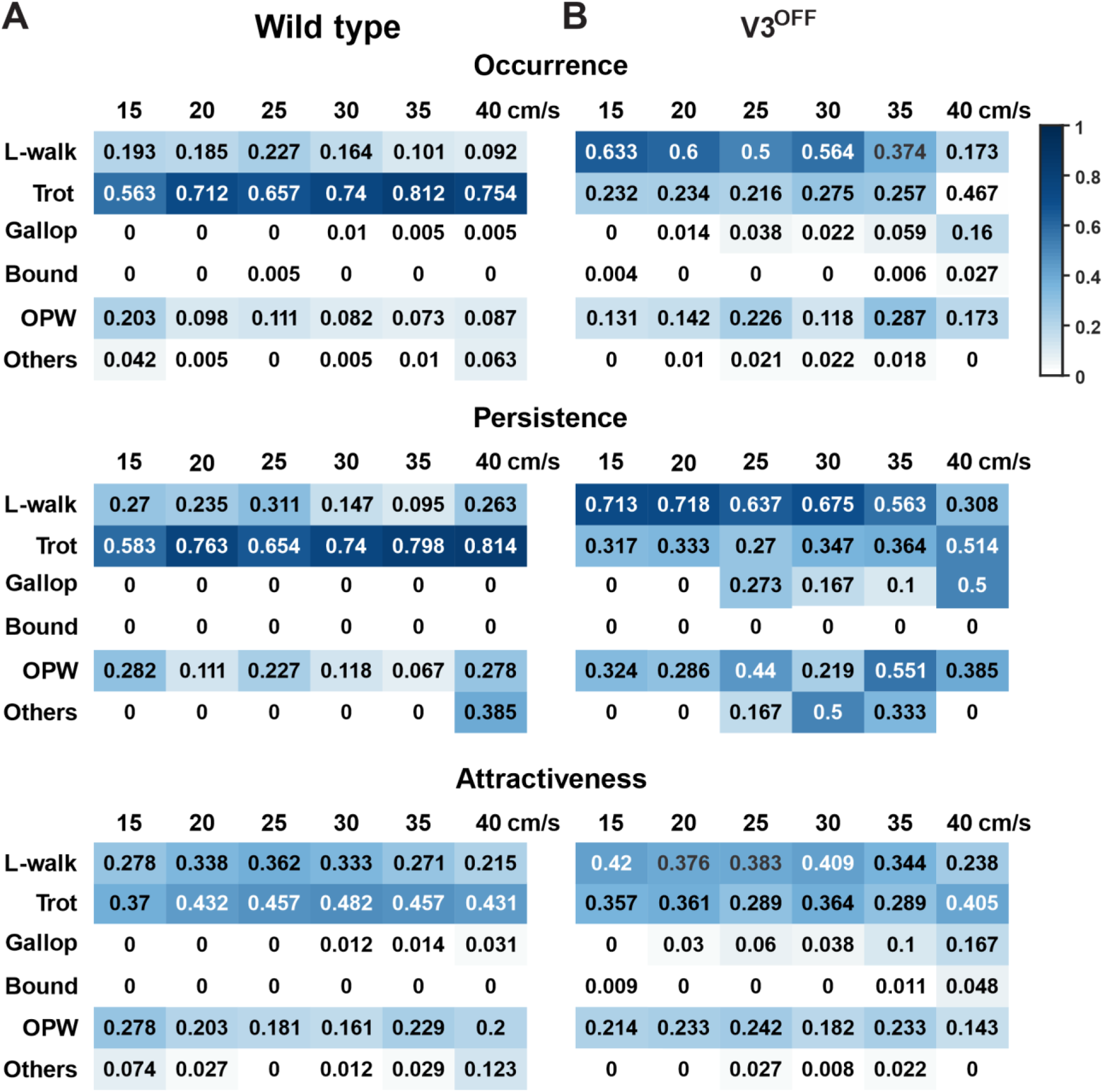
Gait preference of WT and V3OFF mice at individual speeds. Color-coded matrices of occurrence, persistence, and attractiveness values of each type of gait (row) at each tested speed (column). The matrices in **A** are from WT and **B** are from V3^OFF^ mice. The number in each cell shows the value of the parameter of corresponding gait and speed. L-walk: lateral-sequence walk. OPW: out-of-phase walk.

